# Striatal Acetylcholine Dip-Rebound Is Induced by Direct-Pathway Neurons and Encode Action-Outcome Contingency

**DOI:** 10.64898/2026.01.13.699373

**Authors:** Ruifeng Chen, Xueyi Xie, Himanshu Gangal, Xuehua Wang, Jun Wang

**Affiliations:** Department of Neuroscience and Experimental Therapeutics, College of Medicine, Texas A&M University Health Science Center; Institute for Neuroscience, Texas A&M University; Interdisciplinary Faculty of Toxicology, Texas A&M University

## Abstract

Learning contingencies between actions and outcomes is pivotal in adaptive behavior and requires ongoing flexible sensory, motor, and reward information integration. The striatum is central to the integration. However, it remains unclear how the striatal neurons interact with each other to facilitate learning. Here we show that in the dorsomedial striatum, direct-pathway medium spiny neurons (dMSNs), but not dopamine (DA), inhibit cholinergic interneurons, inducing a characteristic acetylcholine (ACh) dip–rebound, encoding action-outcome (A-O) contingency. Using genetically encoded sensors and in vivo fiber photometry, we find that dMSN activation, ACh dip–rebound, and DA transients emerge only after mice acquire the contingency. dMSN activity and ACh dynamics persist even when rewards are probabilistic, adapt in time as the learned relationship between action and reward evolves, and vanish when the contingency is degraded. Ex vivo recordings and in vivo optogenetics further show that dMSN activity is both sufficient and required to generate the ACh dip–rebound through GABAergic inhibition of cholinergic interneurons. Disrupting the dip–rebound slows acquisition and accelerates extinction. Together, these findings reveal a previously unrecognized dMSN–ACh circuit mechanism that encodes contingency during instrumental learning, operating alongside DA signals that track reward outcomes.

## Introduction

Learning contingencies between actions and outcomes is fundamental for survival. These A-O contingencies allow animals to assign motivational value to behaviors that reliably lead to rewards such as food or other resources, thereby shaping future decisions and reinforcing adaptive strategies(Dumas 2005, Fanselow and Poulos 2005, Suzuki 2008, Dickinson 2012, Rosas, Todd et al. 2013, Hawkins and Byrne 2015). Reward-associated actions can become motivational, acquiring incentive salience or ‘wanting,’ which can drive seeking behaviors with learning contingencies. Notably, extensive evidence from both human and animal studies indicates that abnormalities in the learning process contribute to endophenotypes seen in addiction, schizophrenia, depression, Parkinson’s disease, and other psychopathological conditions(Bevins and Palmatier 2004, Hall, Romaniuk et al. 2009, Giovanniello, Bravo-Rivera et al. 2023).

The striatum is a central hub for integrating cortical and thalamic inputs to guide action selection and instrumental learning(Everitt and Robbins 2005, Surmeier 2013, Cox and Witten 2019, Stuber 2023, Jang, Ward et al. 2024). The striatum primarily consists of GABAergic projection neurons known as medium spiny neurons (MSNs), which form two parallel pathways with distinct functions: direct-pathway MSNs(dMSNs), which express dopamine D1 receptors and promote movement and learning through inhibitory projections to basal ganglia output nuclei, and indirect-pathway MSNs(iMSNs), which express D2 receptors and suppress competing actions(Gerfen, Engber et al. 1990, Stuber 2023). Striatal DA, a major neuromodulator of MSNs, plays a critical role in signaling reward prediction errors and in shaping approach–avoidance strategies(Cohen, Haesler et al. 2012, Lerner, Shilyansky et al. 2015, Coddington and Dudman 2018). The striatum is also enriched in acetylcholine (ACh), provided predominantly by cholinergic interneurons (CINs), which have long been theorized to modulate learning and flexibility(Lavoie, Smith et al. 1989, Holt, Graybiel et al. 1997, Morris, Arkadir et al. 2004). Several studies have suggested an equally important role of striatal ACh as DA in cognitive function and motor learning(Woolf and Butcher 1981, Contant, Umbriaco et al. 1996). For example, reward-predicting cues or motivated behavior reliably induced a characteristic ‘pause’ in CIN firing(Aosaki, Tsubokawa et al. 1994, Morris, Arkadir et al. 2004, Skirzewski, Princz-Lebel et al. 2022). Silencing cocaine-induced CIN activity blocked cocaine conditioning(Witten, Lin et al. 2010). However, unlike DA, which is widely accepted to encode reward prediction error, the precise role of ACh remains less clearly defined(Morris, Arkadir et al. 2004, Joshua, Adler et al. 2008, Apicella, Ravel et al. 2011).

To investigate the functional role of striatal neurons and neurotransmitters in learning A-O contingency, we used genetically encoded sensors and fiber photometry to record the dynamics of DA, ACh, and calcium in dMSNs and iMSNs in the dorsal medial striatum during instrumental learning, by which A-O contingencies are formed and strengthened. First, we found that DA, ACh, dMSNs, and iMSNs have distinct responses in well-trained mice in instrumental learning. By systematically uncoupling actions from their outcomes using probabilistic, temporally delayed, and contingency-degradation schedules, we discovered that DA reliably tracks reward delivery, whereas ACh and dMSNs track the contingency between the action and the outcome. Second, we combined ex vivo whole-cell recordings, in vivo optogenetic manipulation, and ACh photometry, showing that dMSN activation is both sufficient and required to generate the ACh dip–rebound via GABAergic inhibition of CINs. Finally, disrupting the ACh dip–rebound during behavior selectively slowed the acquisition of A–O contingencies and accelerated extinction, demonstrating that this dMSN-driven ACh dynamic is essential for establishing and maintaining the contingency.

These findings identify a previously unrecognized dMSN–ACh circuit mechanism that encodes A–O contingency during learning, operating in parallel with DA signals that report reward outcomes. This work revises the classical view of ACh–DA interactions and provides a mechanistic framework for understanding how the striatum contributes to action selection and instrumental learning.

## Results

### A–O contingency formation drives dMSN activity, ACh dip-rebound, and DA release

To elucidate the roles of ACh, DA, and the two major populations of MSNs (dMSNs and iMSNs) in instrumental learning, defined as the establishment of A–O contingencies(Yin and Knowlton 2006), we monitored neurotransmitter dynamics and neuronal activity in the DMS of mice during an operant food self-administration task. We recorded DA and ACh signals in free-moving wild-type mice injected with AAVs encoding GRAB_gDA3h_ and GRAB_gACh4m_(Huang, Chen et al. 2024), respectively. In parallel, D1-Cre and A2A-Cre mice were injected with AAVs-Flex-GCaMP7f to selectively measure dMSN and iMSN activity (sFig. 1). Optical fibers were implanted above the DMS injection sites. After two weeks of recovery, mice were trained on a Fixed Ratio 1 (FR1) schedule(see Methods), in which each active lever press (action) immediately triggered delivery of a food pellet (outcome) (Fig. 1A, 1B).

**Figure 1.**
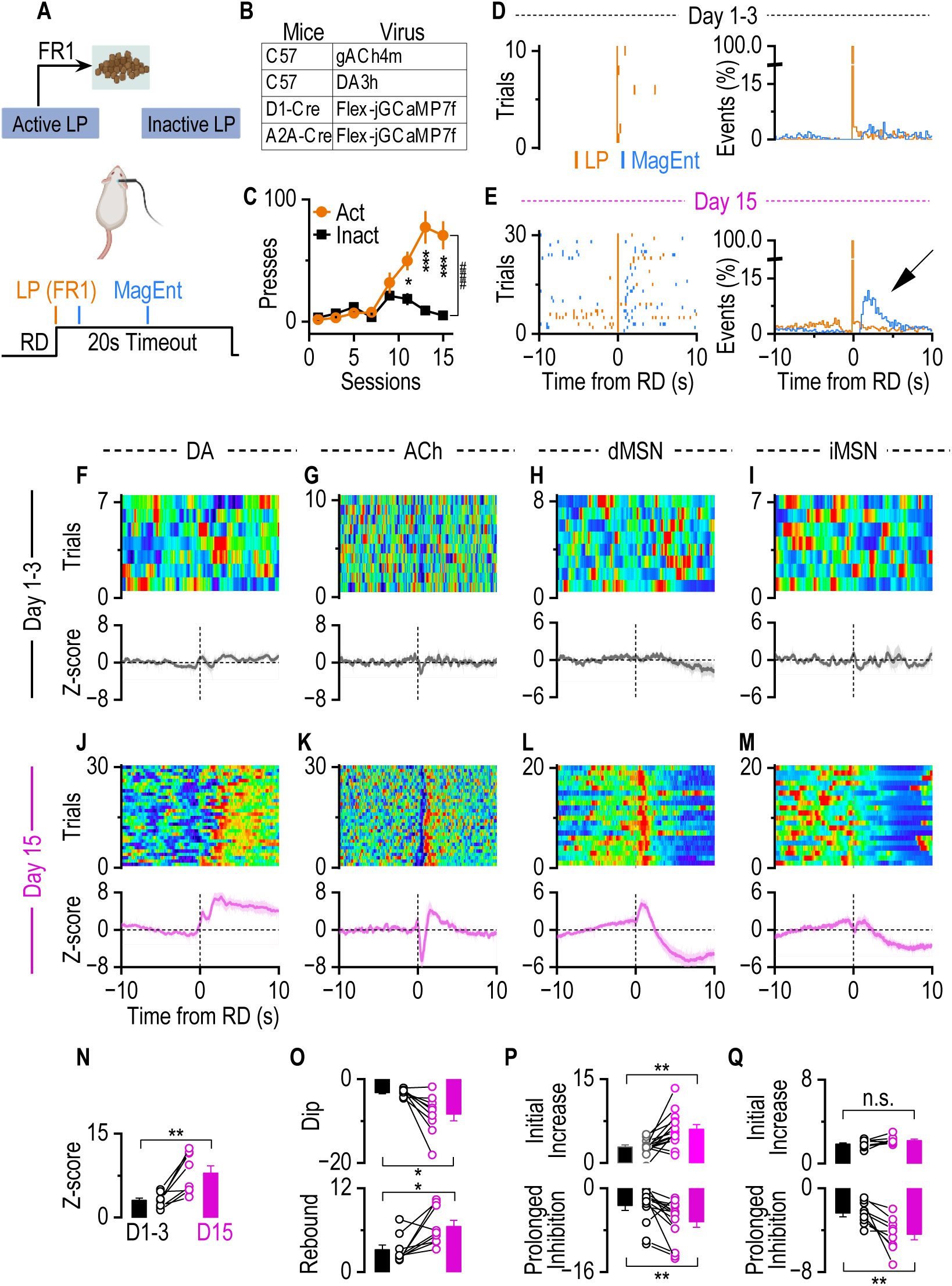
Instrumental learning induced the distinct responses of DA, ACh, dMSNs, and iMSNs after reward delivery. **A.** Schematic representation of the experimental setup. Mice were trained in an operant chamber to press a lever (Active) for triggering reward delivery (RD) using a fixed ratio 1 (FR1) schedule. LP, lever press. **B.** Animals and virus infusion. C57BL/6 mice received bilateral infusion of an AAV-_g_ACh4m or AAV-gDA3h into the DMS. D1-Cre mice or A2A-Cre mice were infused with AAV-Flex-jGCaMP7f into the DMS. **C.** Time course of lever pressing across 15 days, showing increased active presses while inactive presses remained low. ^###^*p* < 0.001 (*F*_(1, 216)_ = 36.22) by 2-way ANOVA with mixed effect analysis; \**p* < 0.05, \*\*\**p* < 0.001 active versus inactive at the same sessions by Sidak post-hoc analysis; n = 14 mice. **D.** Sample Raster plot of behavior trace showing LP and MagEnt aligned to RD from one mouse during the first 3 days (left). The average distribution of LP and MagEnt in all mice (right). n=14 mice. **E.** Sample Raster plot of behavior trace showing LP and MagEnt on the 15^th^ day from one mouse (left). The average distribution of LP and MagEnt for all mice (right, n = 14 mice). Note that there is an increase in magazine entries immediately after LP (indicated by the arrow) on day 15 as compared to days 1-3. **F-I.** No significant DA dynamics, ACh dynamics, and MSN activity are detected during the early phase of instrumental learning. Top, representative heatmap from one mouse across the first 3 days of training. Bottom, averaged signals aligned to lever presses across all mice (DA: n = 8; ACh: n = 9; dMSN: n = 14; iMSN: n = 9 mice). **J-M.** After 15 days of FR1 instrumental training, DA showed an increased dynamic (**J**); ACh showed a dip-rebound dynamics after reward delivery (**K**); dMSN showed an initial increased activity followed by a prolonged inhibition (**L**); iMSN showed a decreased activity without initial activity increase (**M**). Top panels: representative single-mouse heatmaps aligned to RD. Bottom panels: average fiber photometry traces across animals. **N.** DA showed a significant increase after reward delivery in well-trained mice (*t*_(7)_ = 4.39, \*\**p* < 0.01, paired t test). Each pair of linked dots represents one mouse. n = 8 mice. **O.** Instrumental learning enhances both the dip and rebound magnitude of ACh dynamics. Top, magnitude of the dip (*t*_(8)_ = 3.32, \**p* < 0.05, paired t test). Bottom, magnitude of the rebound (*t*_(8)_ = 3.05, \**p* < 0.05, paired t test). n = 9 mice. **P**. dMSNs activity increased significantly after 15 days of training after RD, compared to the first 3 days of training (Top: *t*_(13)_ = 3.73, \*\**p* < 0.01, paired t test). A prolonged inhibition followed the initial activity increase after 15 days of training (Bottom: *t*_(13)_ = 3.14, \*\**p* < 0.01, paired t test). n = 14 mice. **Q.** After 15 days of training, there is a prolonged iMSN inhibition after RD without initial iMSN activity increase (Initial increase, Day 1-3 vs Day 15: *t*_(8)_ = 2.0, *p* > 0.05, paired t test; Prolonged inhibition, Day 1-3 vs Day 15: *t*_(8)_ = 3.64, \*\**p* < 0.01, paired t test). n = 9 mice.

Behavioral analysis showed that mice successfully differentiated which lever was associated with reward over 15 sessions, evidenced by a significant increase in active relative to inactive lever presses (Fig. 1C). Early in training, active lever presses were not consistently followed by magazine entries, indicating exploratory behavior and absence of an established contingency (Fig. 1D). By day 15, lever presses were reliably followed by immediate magazine entries, confirming a learned association between the action and outcome (Fig. 1E). Fiber photometry recordings mirrored the behavioral progression of learning. During the first three training sessions, reward delivery following lever presses evoked minimal changes in DA and ACh dynamics, as well as limited activity in both dMSNs and iMSNs (Fig. 1F-1I). Magazine entries also produced little response (sFig. 3), indicating that neither the reward-seeking (lever press) nor reward-taking (magazine entry) behaviors themselves were sufficient to elicit neuronal or neurotransmitter responses in the DMS. After 15 sessions, however, clear and consistent activity patterns emerged. Coinciding with increased DA levels (Fig. 1J), ACh release displayed a characteristic dip–rebound pattern aligned to reward delivery (Fig. 1K), and dMSN activity displays an initial increase followed by a prolonged inhibition (Fig. 1L). iMSNs showed transient activation after reward delivery during early learning, which diminished with continued training (sFig. 4). Interestingly, iMSN, similar to dMSN, exhibits a prolonged suppression phase, but without initial activity increase in well-trained mice (Fig. 1M). Quantitative analysis revealed that 15 sessions of training significantly enhanced the DA transient after reward delivery (Fig. 1N). The ACh dip and rebound amplitudes were significantly increased in the last session compared to the first three sessions. On day 15, the dip occurred at 0.59 ± 0.06 sec, and the rebound occurred at 1.72 ± 0.24 sec after reward delivery (Fig. 1O). Consistent with these changes, the initial dMSN activity increase and the prolonged inhibition were elevated after reward delivery on day 15 compared to early sessions (Fig. 1P). iMSNs showed a prolonged inhibition without an immediate increase following reward delivery (Fig. 1Q).

**Figure 2.**
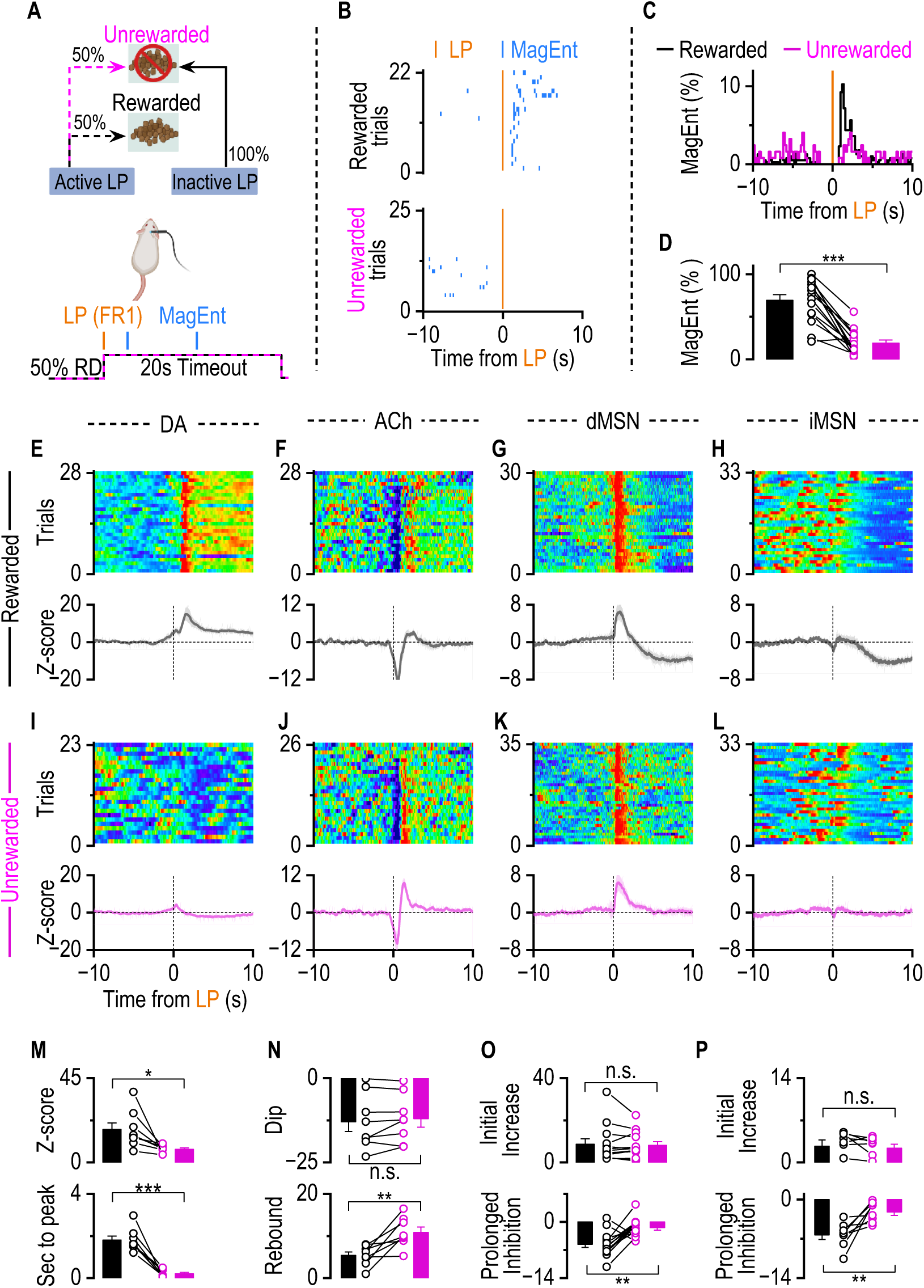
DA Release Tracks Reward, whereas dMSNs and ACh do not. **A.** Schematics of a schedule in which an active lever press (LP) is followed by either a reward (with a 50% chance) or a 10s gap without reward. There is a fixed ratio 1 (FR1) schedule where one lever press leads to a reward or pseudo-reward. **B.** Raster plot of sample behavior trace of rewarded trials (top) and unrewarded trials(bottom) from a mouse showing LP(Orange) and MagEnt(Blue) aligned to LP. **C.** Average distribution of MagEnt before and after LP. n = 14 mice. **D.** Summary data comparing the percentage of MagEnt immediate (<3s) after LP, where mice enter the magazine during rewarded trials versus unrewarded trials. rewarded trials show high immediate magazine entry (black bar), while unrewarded trials show significantly less immediate magazine entry (pink bar, *t*_(13)_ = 7.29, \*\*\**p* <0.001, paired t test). n = 14 mice. **E-H.** In rewarded trials, DA, ACh, dMSNs, and iMSNs showed similar dynamics as in Fig. 1f-DA showed an increased dynamic (**E**); ACh showed a dip-rebound dynamics after reward delivery (**F**); dMSN showed an increased activity followed by a decreased activity (**G**); iMSN showed a decreased activity without initial activity increase (**H**). Top panels: representative single-mouse heatmaps aligned to LP. Bottom panels: average fiber photometry traces across animals (DA: n = 8; ACh: n = 8; dMSN: n = 14; iMSN: n = 8). **I-L.** In unrewarded trials, DA dynamics, ACh dynamics, dMSNs, and iMSNs activity showed different patterns as in Figure 2e-h. ACh showed a dip-rebound dynamics after reward delivery, while the rebound is significantly bigger than that in reward trials (**E**); DA showed a less significant increase (**F**); dMSN showed an increased activity without prolonged inhibition following (**G**); iMSN showed no significant response (**H**). Top panels: representative single-mouse heatmaps aligned to LP from the same mouse in panels **E-H**. Bottom panels: average fiber photometry traces across animals. **M.** Comparison of the DA release between rewarded trials and unrewarded trials. The amplitude of DA release in RD trials is significantly higher than that in unrewarded trials(*t*_(7)_ = 3.36, \**p* <0.05, paired t test), and the peak latency was significantly longer in rewarded trials(*t*_(7)_ = 5.66, \*\*\**p* <0.001, paired t test). n = 8 mice. **N.** The amplitude of the dip (left) and rebound (right) of Z-scored ACh signals in rewarded and unrewarded trials. The dip amplitudes were comparable between conditions(*t*_(7)_ = 1.16, *p* >0.05, paired t test), while the rebound amplitude was significantly larger in unrewarded trials (*t*_(7)_ = 3.80, \*\**p* <0.01, paired t test). n = 8 mice. **O.** No significant difference in the peak intensity of dMSN initial activity between rewarded and unrewarded trials(*t*_(13)_ = 0.69, *p* >0.05, paired t test). This suggests that dMSN activation patterns are stable across different trial outcomes, indicating a consistent dMSN response irrespective of reward outcome. While the prolonged inhibition following the initial activity increase was diminished in the unrewarded trials (*t*_(13)_ = 3.78, \*\**p* <0.01, paired t test). **P.** iMSNs activity showed a prolonged inhibition without an initial activity increase in rewarded trials after RD. In unrewarded trials, the prolonged inhibition was diminished (Initial increase: rewarded trials vs. unrewarded trials, *t*_(7)_ = 0.48, *p* >0.05, paired t test; prolonged inhibition: rewarded trials vs. unrewarded trials, *t*_(7)_ = 3.44, \**p* <0.05, paired t test).

**Figure 3.**
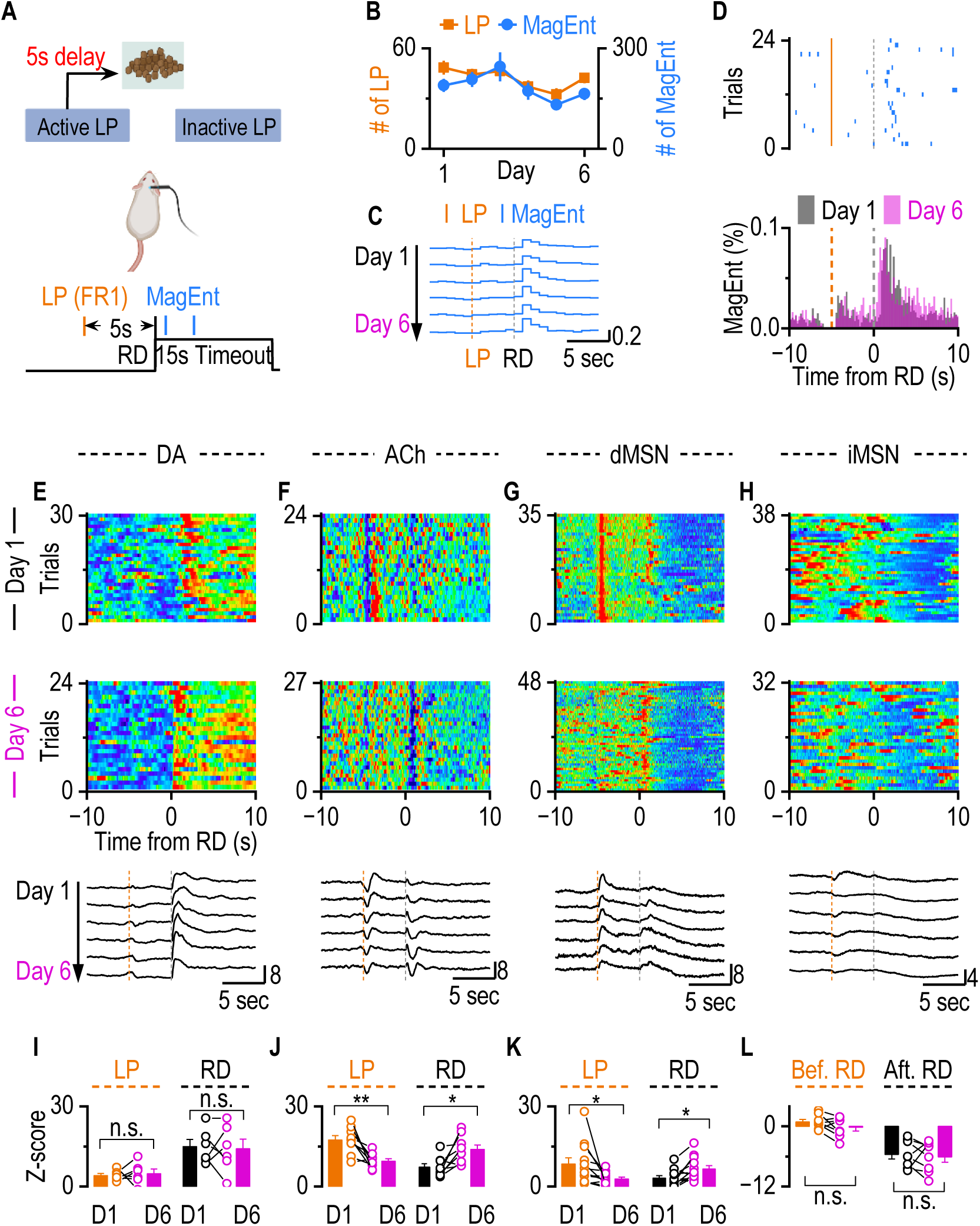
Temporal Shifts in dMSN Activity and ACh Dip-Rebound from LP to RD During Immediate Association Decoupling Training. **A.** We adopt a Fixed Ratio with Fixed Delay (FR1-FD5) Schedule after the VR2 schedule with the same mice, where each LP is followed by a 5-second delay before reward delivery. LP will induce lever withdrawal until the end of the reward delivery period, preventing the initiation of the next trial. **B.** There is no difference between the number of LP and MagEnt on day 1 and day 6 (LP: day 1 vs. day 6, *t*_(13)_ = 1.48, *p* =0.16, paired t test; MagEnt: day 1 vs. day 6, *t*_(13)_ = 1.26, *p* =0.23, paired t test). **C-D.** The distribution of MagEnt during immediate association decoupling training. **C.** The distribution of MagEnt from day 1 to day 6. **D.** Sample raster plot of MagEnt from one mouse in one session (Top). Orange lines represent individual lever presses. Blue ticks represent magazine entries, which indicate when the mouse enters the magazine to check for a reward. The x-axis is aligned around the time of reward delivery, with time zero marking reward delivery. Average MagEnt distribution across all mice on day 1 and day 6(Bottom). **E.** DA transients consistently occurred after RD on both Day 1 and Day 6, without evidence of a temporal shift. Top: Sample heatmap of DA dynamics aligned to RD from one mouse on Day 1. Middle: Sample heatmap from the same mouse on Day 6. Bottom: Averaged DA dynamics across all mice from Day 1 to Day 6, aligned to RD, showing stable post-reward increases in DA release. **F.** Temporal shift of ACh dip-rebound dynamics during immediate association decoupling training. On Day 1, ACh dip-rebound occurred after lever press (LP) but not after reward delivery (RD). Across six days of training, the dip-rebound gradually shifted from LP to RD. Top: Sample heatmap of ACh dynamics aligned to RD from one mouse on Day 1. Middle: Sample heatmap from the same mouse on Day 6. Bottom: Averaged ACh dynamics across all mice from Day 1 to Day 6, aligned to RD, showing the progressive shift of the ACh dip-rebound. **G.** Consistent with ACh dynamics, dMSN activity showed a Temporal shift from LP to RD during immediate association decoupling training. On Day 1, dMSN transients occurred after lever press (LP) but not significantly after reward delivery (RD). Across six days of training, the transient gradually shifted from LP to RD. Top: Sample heatmap of dMSN activity aligned to RD from one mouse on Day 1. Middle: Sample heatmap from the same mouse on Day 6. Bottom: Averaged dMSN activity across all mice from Day 1 to Day 6, aligned to RD, showing the progressive shift of the dMSN transient toward RD. **H.** iMSN shows no immediate response following either LP or RD. Top: Sample heatmap of iMSN activity aligned to RD from one mouse on Day 1. Middle: Sample heatmap from the same mouse on Day 6. Bottom: Averaged iMSN activity across all mice from Day 1 to Day 6, aligned to RD. **I.** Showing the DA release over time for both RD and LP on Day 1 and Day 6(LP: day 1 vs. day 6, *t*_(5)_ = 0.32, *p* =0.77, paired t test; RD: day 1 vs. day 6, *t*_(5)_ = 0.22, *p* =0.83, paired t test). The amplitude of DA transients following either LP or RD did not change over time. **J.** Comparison of the ACh dip-rebound magnitude on Day 1 versus Day 6 after LP and RD (LP: day 1 vs. day 6, *t*_(8)_ = 3.86, \*\**p* <0.01, paired t test; RD: day 1 vs. day 6, *t*_(8)_ = 3.19, \**p* <0.05, paired t test). ACh dip-rebound shifts from being associated with lever press early in training to becoming more linked with reward delivery as learning progresses. **K.** Comparison of the dMSN activity on Day 1 versus Day 6 after LP and RD (LP: day 1 vs. day 6, *t*_(11)_ = 2.42, \**p* <0.05, paired t test; RD: day 1 vs. day 6, *t*_(11)_ = 2.96, \**p* <0.05, paired t test). dMSNs activity following events shifts from being associated with lever press early in training to becoming more linked with reward delivery as learning progresses. **L.** There is no activity increase before RD across 6 days of training (Day 1 vs Day 6: *t*_(7)_ = 1.69, *p* >0.05, paired t test). The prolonged inhibition after RD didn’t change across 6 days of training (Day 1 vs. Day 6: *t*_(7)_ = 0.46, *p* >0.05, paired t test).

**Figure 4.**
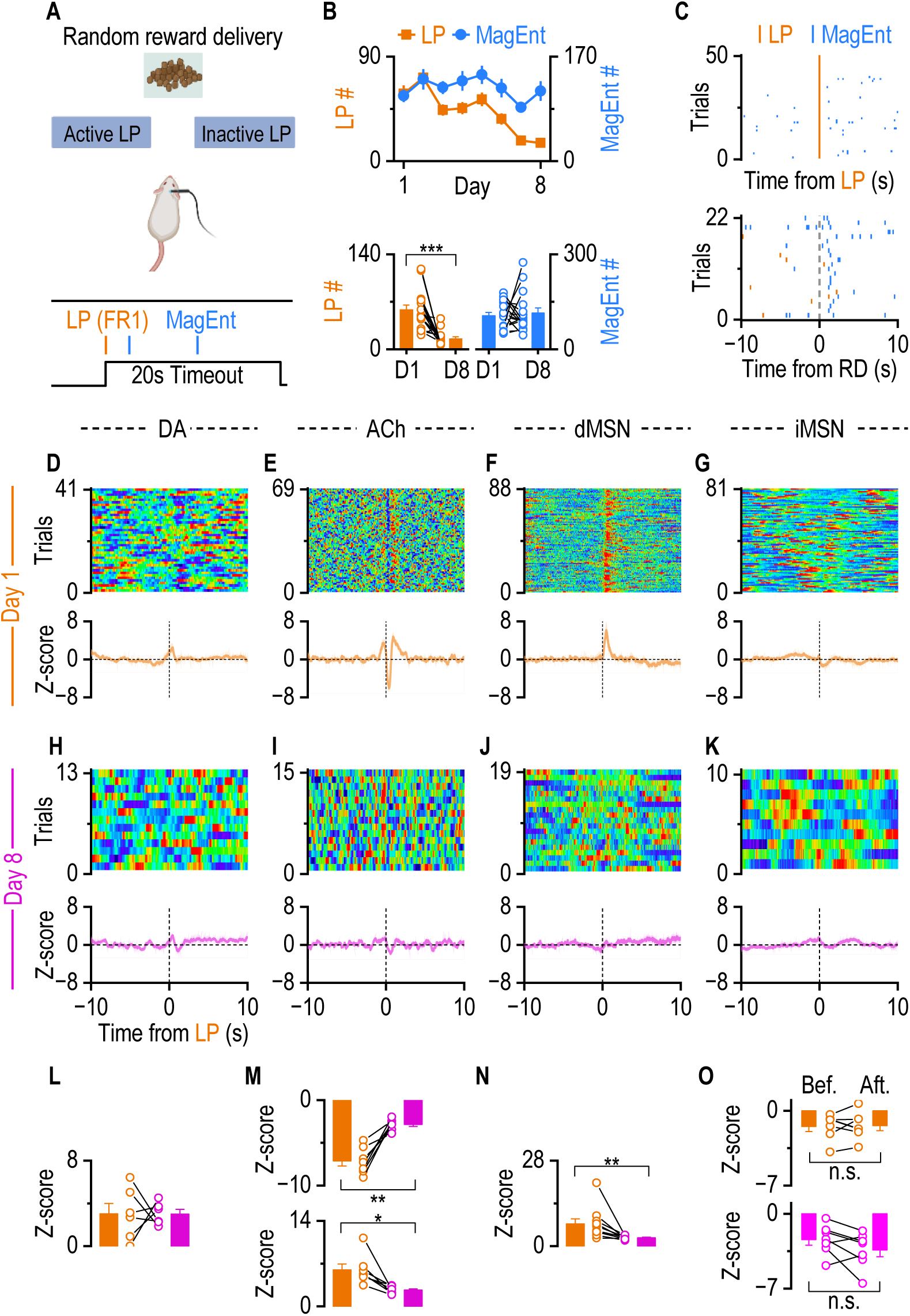
Contingency Degradation Training Eliminates the Dip-Rebound in ACh Release and dMSN Activity. **A.** We adopt a contingency degradation behavior schedule after Immediate Association Decoupling Training with the same mice, where LP initiates a 10s gap without RD. The food reward was delivered at a variable interval (on average 80s), independent of LP. With the schedule, LP is not associated with reward delivery, so the action-outcome contingency was broken. **B.** The number of LP and MagEnt from day 1 and day 8. The number of LP significantly decreased from day 1 to day 8 (*F* = 21.79, \*\*\**p* < 0.001, RM-ANOVA). While there is no difference in the number of MagEnt between the first day and the last day (*F* = 2.29, *p* = 0.082, RM-ANOVA). **C.** Sample raster plot shows immediate MagEnt follows RD but not LP. **D-G.** On the first day of contingency degradation training, ACh and dMSN maintained the response to LP. DA showed a tiny increase initiated before LP (**D**). ACh showed a dip-rebound following an initial burst (**E**); dMSN showed an increased activity after LP (**F**); iMSN showed no response to LP (**G**). Top panels: representative single-mouse heatmaps aligned to LP. Bottom panels: average fiber photometry traces across animals (DA: n = 6; ACh: n = 7; dMSN: n = 10; iMSN: n = 7). **H-K.** After 8 days of contingency degradation training, ACh and dMSN lost the response to LP. ACh(**H**), DA(**I**), dMSN(**J**), and iMSN(**K**) showed almost no response to LP. Top panels: representative single-mouse heatmaps aligned to LP. Bottom panels: average fiber photometry traces across animals. **L.** Statistics show that the amplitude of DA response was not significantly changed from day 1 to day 8 (*t*_(5)_ = 0.009, *p* =0.99, paired t test). **M.** Statistics show that the amplitude of ACh dip and rebound was significantly diminished from day 1 to day 8 (Dip: day 1 vs. day 8, *t*_(6)_ = 5.87, \*\**p* <0.01, paired t test; rebound: day 1 vs. day 8, *t*_(6)_ = 3.61, \**p* <0.05, paired t test). **N.** dMSN response after LP was significantly reduced from Day 1 to Day 8 (*t*_(9)_ = 2.85, \*\**p* <0.01, paired t test). **O.** There is no difference in iMSN activity before and after LP on Day 1 and Day 8 (Day 1: before LP vs. after LP, *t*_(6)_ = 0.26, *p* >0.05, paired t test; Day 8: before LP vs. after LP, *t*_(6)_ = 1.74, *p* >0.05, paired t test).

Together, these findings suggest that the acquisition of instrumental learning, rather than performing the exploratory action itself, evokes striatal dMSN activity, the ACh dip-rebound, and DA phasic release.

### DA tracks reward, whereas dMSNs and ACh do not

Instrumental learning includes action, reward, and the contingency between action and outcome. Because the FR1 schedule couples the reward-related action (lever press) and the guaranteed outcome (food pellet) closely in time, it cannot untangle their individual contributions. To identify the specific roles of ACh, DA, and MSNs, we next dissociated action from outcome using a variable ratio (VR2) schedule (see Methods), employing the same animals previously trained under the FR1 condition in Figure 1. Under this VR2 schedule, each active lever press resulted in a 50% probability of reward, creating two distinct trial types: rewarded trials and unrewarded trials (Fig. 2A). In rewarded trials, lever pressing resulted in food delivery and a 20-second timeout. Unrewarded trials involved a lever press followed by a 20-second timeout without reward delivery. Behavioral analysis revealed clear distinctions between trial types: mice consistently exhibited immediate magazine entry following lever presses in rewarded trials with magazine entry peak at 1.5 sec post-reward delivery (Fig. 2B, consistent with Fig. 1E), but rarely entered the magazine immediately in unrewarded trials (Fig. 2C). Quantitative analysis confirmed this observation, showing significantly fewer immediate magazine entries (defined as magazine entries within 3 sec post lever press divided by the number of trials) in unrewarded trials compared to rewarded trials (Fig. 2D). The absence of immediate magazine entries in unrewarded trials confirmed that the mice detected the lack of reward in unrewarded trials.

Fiber photometry recordings demonstrated distinct neurotransmitter dynamics and MSN activity in rewarded versus unrewarded trials. In rewarded trials, lever presses triggered a marked DA rise, a characteristic ACh dip-rebound pattern, and an initial increase in dMSN activity, accompanied by unaltered iMSN activity (Fig. 2E-2H, consistent with Fig. 1J-1M). In contrast, unrewarded trials elicit minimal DA rise, while the ACh dip-rebound pattern and initial MSN activity remain similar in rewarded trials, except that the prolonged inhibition of MSN disappeared (Fig. 2I-2L).

Further statistical analyses showed that the magnitude and timing of peak DA rise were substantially reduced in unrewarded versus rewarded trials (Fig. 2M). The magnitude of ACh dip did not differ between rewarded trials and unrewarded delivery trials (Fig. 2N top); however, the ACh rebound was significantly greater in unrewarded versus rewarded trials (Fig. 2N bottom). The decreased ACh rebound observed in reward trials (Fig. 2N) may be due to enhanced DA release triggered by reward, as DA can inhibit CINs via DA D2 receptors (Straub, Tritsch et al. 2014, Martyniuk, Torres-Herraez et al. 2022, Chantranupong, Beron et al. 2023). dMSNs consistently exhibited similar increased activity in both rewarded and unrewarded trials, implying that the initial dMSN activity may encode the reward-related action or the contingency rather than the outcome (Fig. 2O top). Again, iMSNs did not show immediate significant changes in activity after lever pressing (Fig. 2P top). The prolonged MSN inhibition in rewarded trials disappeared in unrewarded trials (Fig. 2O bottom, Fig. 2P bottom).

Together, these results suggest ACh and DA have distinct functional roles in instrumental learning. DA primarily tracks reward outcomes, consistent with its well-established role in encoding reward prediction errors (Schultz, Dayan et al. 1997, Waelti, Dickinson et al. 2001, Steinberg, Keiflin et al. 2013, Amo 2024). In contrast, ACh dip-rebound and dMSN activity still exist in the absence of reward, although the magnitude of ACh rebound is modified.

### dMSNs and ACh Track A-O Contingency

Depends on the observations from Figure 2, we hypothesized that dMSN activity and ACh dynamics represent the reward-related action rather than the exploratory action or reward itself. To test this hypothesis, we employed a Fixed Ratio 1 with Fixed Delay 5-second schedule (FR1-FD5) in which a 5-second delay was introduced between lever press and reward delivery, to temporally dissociate the action from its outcome (Fig. 3A), allowing us to determine whether dMSN activation and ACh dip-rebound are aligned with the reward-related action.

The same cohort of mice previously used in Figure 2 were retained using the FR1 schedule (Fig. 1) before the FR1-FD5 training. Across six consecutive sessions of FR1-FD5 training, the total number of lever presses and magazine entries remained stable (Fig. 3B). Magazine entries consistently occurred immediately following reward delivery rather than following the lever press across all training sessions (Fig. 3C, 3D), implying the behavior engagement and performance were unchanged from day 1 to day 6 of the FR1-FD5 training. Fiber photometry recordings revealed DA release increased significantly only after reward delivery, but minimally after lever pressing across 6 days of the training (Fig. 3E). In contrast, increased dMSN activity and a robust ACh dip–rebound pattern immediately following the lever press, but not at the onset of reward delivery on the first day of FR1-FD5 training (Fig. 3F, 3G top). These findings are consistent with our previous expectation that dMSNs activity and ACh dip-rebound track reward-related action rather than outcomes, while DA tracks outcomes. To our surprise, after six days of training, the ACh dip-rebound and dMSN activity were observed both after reward delivery and after lever pressing, indicating a temporal shift in neural activity (Fig. 3F, 3G middle). This shift was gradual, becoming progressively more pronounced across training days (Fig. 3F, 3G bottom). The signal shifts to the proximal predictor, implying an evolving representation. iMSN did not show activity increase after either lever presses or reward deliveries (Fig. 3H). Quantitative analyses confirmed that DA signal consistently maintains high following reward delivery but did not significantly change between day 1 and day 6, remaining stable and specifically associated with reward delivery (Fig. 3I). In contrast, by day 6, the magnitude of the ACh dip-rebound significantly decreased following lever presses but significantly increased following reward delivery, compared to day 1 (Fig. 3J). Similar to ACh dynamics, the dMSN activity following lever presses showed a decrease, whereas the activity after reward delivery showed an increase over the course of training (Fig. 3K). iMSNs consistently exhibited no immediate activity increase after lever pressing but displayed a significant reduction in firing after reward delivery (Fig. 3L).

Taken together, even the behavioral performance from day 1 to day 6 was unchanged, the ACh dynamics and dMSN activity were temporally shifted from lever presses to reward delivery. At the introduction of the 5-s delay, dMSN and CIN activity were coupled to the lever-press, which predicted reward delivery depending on the previous training history. As learning progressed, this activity shifted toward the reward-delivery epoch, suggesting that dMSN and ACh signaling adapt the evolving action–outcome contingency, not just the reward-related action.

### Contingency Degradation Eliminates Post-action dMSN Activity and ACh Dip-Rebound

Wrap up all the results we have so far, dMSN activity and ACh dip-rebound track neither the action nor the outcome in instrumental learning. They are potentially tracking the contingency, the bridge between the action and the outcome. To test the hypothesis that dMSN activity and the ACh dip-rebound pattern represent contingency, a contingency degradation schedule(Liljeholm, Tricomi et al. 2011, Fanelli, Klein et al. 2013, Shillinglaw, Everitt et al. 2014) was implemented with previously trained mice. In this schedule, each lever press no longer results in food delivery. Instead, food was delivered randomly at an average interval of 80 sec (Fixed Ratio with Variable Interval 80, FR1-VI80, See Method; Fig. 4A).

Under this contingency-degraded schedule, the lever press action was dissociated from the reward outcome. After 8 days of training, the number of lever presses significantly decreased compared to day 1, whereas the number of magazine entries remained unchanged (Fig. 4B). This reduction in lever presses indicates that mice recognized and adapted to the degraded contingency between action (lever press) and outcome (reward). Mice consistently entered the magazine immediately following reward delivery rather than after lever presses (Fig. 4C), which demonstrates that the motivation to collect rewards was unaffected. Collectively, these data confirm that the schedule successfully degraded the action-outcome contingency after 8 days of training.

Fiber photometry recordings further supported these behavioral results. On the first day of contingency degradation training, DA shows minimal response to lever presses, but a significant response to the reward delivery, aligning with observations from unrewarded trials and rewarded trials in Figure 2 (Fig. 4D, sFig. 5A). The ACh dip-rebound and dMSN activity were observed following lever presses and reward delivery(Fig. 4E, 4F, sFig. 5B, 5C). While an initial burst was accompanied, potentially due to increased glutamatergic inputs from the cortex or thalamus(Bradfield, Bertran-Gonzalez et al. 2013, Doig, Magill et al. 2014, Krok, Maltese et al. 2023). iMSNs exhibited no significant response to lever presses or reward delivery (Fig. 4G, sFig. 5H). By day 8 of contingency degradation, DA consistently showed minimal response to lever presses, while both the ACh dip-rebound and dMSN activity increase had disappeared after lever presses (Fig. 4H-4J), but remained after reward delivery (sFig. 5E-5G), consistent with expectations, indicating they specifically tracked the contingency. iMSN activities showed no substantial changes between days 1 and 8 (Fig. 4G, 4K, sFig. 5D, 5H). Statistical analysis confirmed that while DA remained statistically unchanged (Fig. 4L), a significant reduction in the ACh dip-rebound and dMSN activity following lever presses across the training period (Fig. 4M, 4N). iMSN shows no response to lever presses (Fig. 4O). These findings collectively demonstrate that contingency degradation effectively abolishes the ACh dip-rebound and dMSN activity response, confirming their role in representing the contingency.

**Figure 5.**
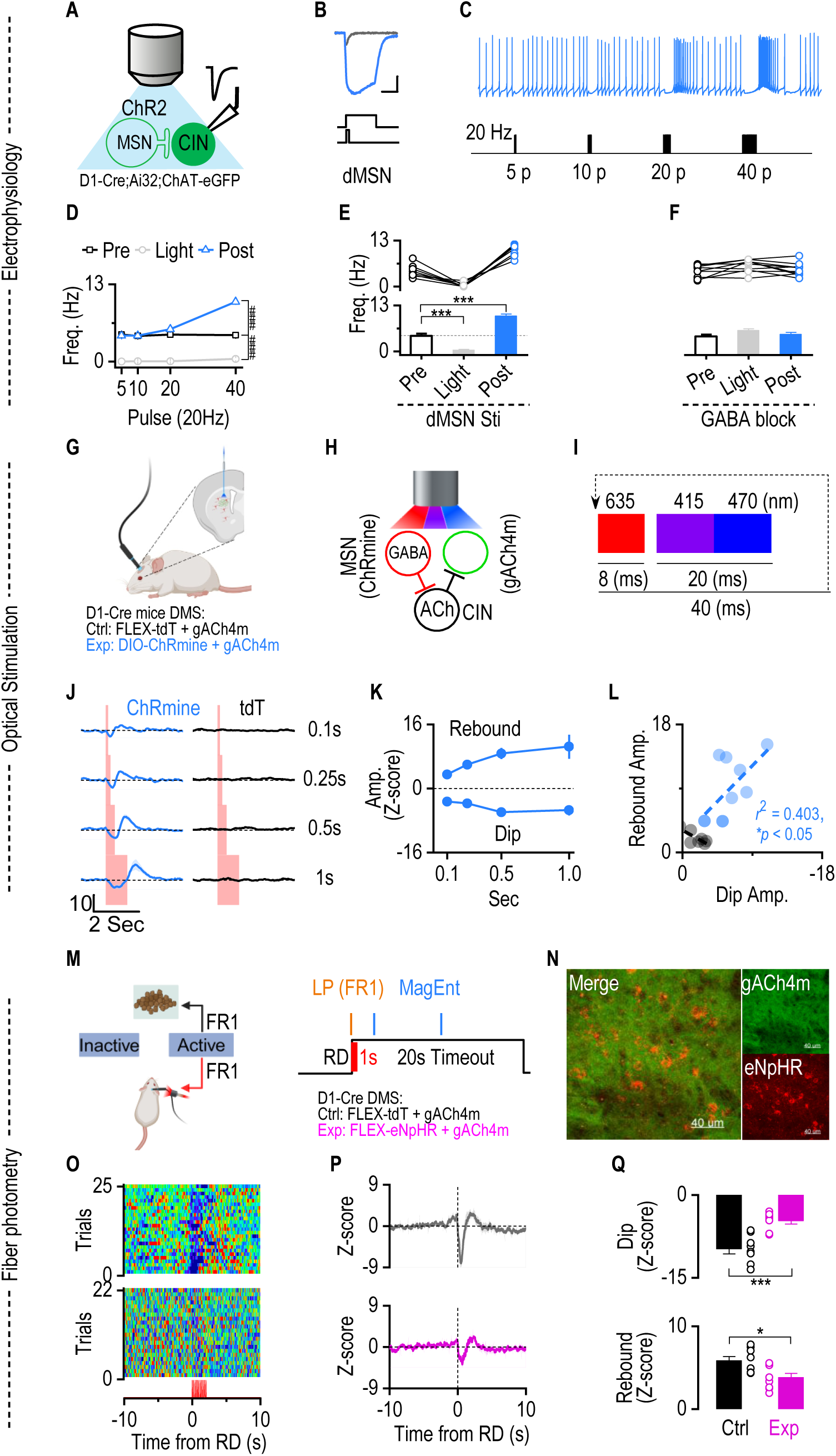
dMSN excitation is sufficient and required to induce ACh dip-rebound in vivo. **A.** Schematic of the experimental setup for optogenetic stimulation of dMSNs expressing ChR2 and whole-cell recordings from eGFP-labeled CINs in D1-Cre; Ai32; ChAT-eGFP mice. **B.** Example traces showing dMSN membrane depolarization in response to short vs. prolonged blue laser stimulation. **C.** Representative whole-cell current-clamp recording showing CIN firing in the response of dMSN stimulation. dMSN stimulation (20 or 40 pulses, 25 Hz) induces an inhibitory effect on CIN firing, followed by a rebound, compared to short stimulation (5 or 10 pulses, 25 Hz). **D.** Frequency of CIN firing to 20 Hz stimulation of dMSN across different pulses. Line graph showing average firing frequency (Hz) of Pre-stimulation (black squares), during light stimulation (gray circles), and Post-stimulation (blue triangles) in response to increasing pulse number (5, 10, 20, 40 pulses) delivered at 20 Hz (n = 12 neurons from 3 mice). Pre vs. During: ^###^*p* < 0.001 (*F*_(1, 48)_ = 15.24) by 2-way ANOVA with mixed effect analysis; Pre vs. Post: ^###^*p* < 0.001 (*F*_(1, 48)_ = 20.82) by 2-way ANOVA with mixed effect analysis **E.** Frequency of CIN firing pre-, during, and post-optogenetic stimulation of dMSN stimulation. Each line represents data from an individual cell (n = 12 from 3 mice). Group data show a significant increase in frequency post-stimulation compared to both pre- and light-stimulation conditions (Pre vs. During, *t*_(7)_ = 5.45, \*\*\**p* <0.001, paired t test; Pre vs. Post, *t*_(7)_ = 12.83, \*\*\**p* <0.001, paired t test). **F.** Bath application of GABA_A_R antagonist Bicuculline blocked the CIN firing pause during the optical stimulation and diminished the rebound post-stimulation. **G.** Diagram of the combined experimental setup for D1-Cre mice, showing simultaneous optogenetic stimulation and fiber photometry to assess ACh release following dMSN activation. In the control group, D1-Cre mice were infused with AAV-FLEX-tdT and AAV-gACh4m. **H.** Simplified circuit diagram depicting GABAergic input from dMSNs onto CINs and the resulting ACh release, monitored by the gACh4m sensor. **I.** Stimulation protocol: A cyclic sequence of three excitation wavelengths was used—635 nm (8 ms) for optogenetic activation of ChRmine, 415 nm (10 ms) as a photometry reference, and 470 nm (10 ms) for excitation of gACh4m. The full cycle lasted 40 ms and was continuously repeated to enable simultaneous optogenetic stimulation and real-time monitoring of ACh dynamics via fiber photometry. **J.** In vivo fiber photometry traces of ACh signals during optogenetic stimulation of dMSNs or control at increasing durations (0.1, 0.25, 0.5, and 1 s). ACh dip and rebound become more pronounced with longer dMSN stimulation (n = 7 (Ctrl) and 10 (Exp) mice). **K.** Quantification of ACh dip and rebound amplitudes evoked by optogenetic stimulation of dMSNs at varying durations. For both cell types, the dip and rebound amplitudes increased with stimulation duration and peaked at 0.5 seconds. **L.** A negative correlation between ACh dip and rebound amplitudes was observed following dMSN stimulation (*r* = -0.63, slope = -1.13). Each dot represents one mouse in response to 25 Hz, 1-second stimulation. **M-N.** Schematic of behavioral setup. D1-Cre mice were infused bilaterally in the DMS with AAVs-GRAB-ACh4m alone (Ctrl) or AAVs-GRAB-ACh4m plus AAV-FLEX-eNpHR3.0 (Exp). An optical fiber implanted into the injection site to enable simultaneous red laser delivery (635 nm, 10 Hz, 1 s) to inhibit dMSNs and blue LED excitation for ACh photometry. Recovered mice underwent an FR1 schedule with a 100% chance of 1-second laser delivery after the initiation of reward delivery. **O-Q.** The 1-second optical inhibition of dMSN activity after RD significantly diminished ACh dip rebound. **O.** Representative single-mouse heatmaps aligned to LP from the Ctrl (Top) and Exp (Bottom) group. **P**. Average fiber photometry traces across animals (Ctrl: n = 8 mice; Exp: n = 8 mice). **Q.** Statistics show that the ACh dip and rebound in the Exp group were less significant than the Ctrl group (Dip: *t*_(14)_ = 4.81, \*\*\**p* < 0.001, unpaired t test; rebound: *t*_(14)_ = 2.75, \**p* < 0.05, unpaired t test).

### dMSN excitation is sufficient and required to induce ACh dip-rebound in vivo

Previous electrophysiological studies have shown that optical stimulation of GABAergic neurons or dopaminergic terminals in the striatum, with acute brain slices, induces a pause-rebound pattern in ACh firing(Aosaki, Graybiel et al. 1994, Brown, Tan et al. 2012, Straub, Tritsch et al. 2014, Dorst, Tokarska et al. 2020, Gangal, Xie et al. 2023). This suggests that both DA release and MSN activity are potential candidates responsible for the ACh dip-rebound observed during training sessions. However, temporal discrepancies between DA elevations and ACh dip-rebound observed in our training experiments challenge the role of DA as the primary driver. Thus, we investigated whether dMSNs selectively modulate this ACh dynamic.

We have previously established the anatomical validity of this circuit, demonstrating a direct, monosynaptic projection from dMSNs to striatal CINs using rabies-mediated retrograde tracing (Gangal, Xie et al. 2023). To determine whether activation of dMSNs is sufficient to induce the pause-rebound in CIN firing, we performed whole-cell current-clamp recordings from eGFP-labeled CINs in acute slices prepared from D1-Cre; Ai32; ChAT-eGFP mice (Fig. 5A). Blue-light stimulation of ChR2-expressing dMSNs evoked reliable inward currents in dMSNs (Fig. 5B). Brief dMSN stimulation (5–10 pulses at 20 Hz) had minimal effect on CIN firing, whereas longer stimulation trains (20–40 pulses at 20 Hz) transiently suppressed CIN spiking, followed by a pronounced rebound increase in firing rate (Fig. 5C-5E). Bath application of the GABA_A_ receptor antagonist bicuculline abolished the light-evoked pause and attenuated the rebound response (Fig. 5F).

We then tested whether optical stimulation of dMSN leads to striatal ACh dip-rebound in vivo using fiber photometry to monitor ACh levels in awake D1-Cre mice. AAV-DIO-ChRmine and AAV-gACh4m were infused into the DMS of D1-Cre mice as the experimental group, allowing selective optical stimulation of dMSNs in vivo, and AAV-FLEX-tdT and AAV-gACh4m were infused into the DMS of D1-Cre mice as the control group. Optical fibers were implanted into the injection sites to simultaneously record ACh dynamics using fiber photometry (Fig. 5G,5H). A stimulation protocol combining three excitation wavelengths—635 nm (8 ms laser) for ChRmine activation, 415 nm (20 ms LED) for reference, and 470 nm (20 ms LED) for GRAB_gACh4m_ excitation—was delivered in a 40-ms cycle to enable real-time ACh monitoring combined with optical stimulation (Fig. 5I). Our results show that optical stimulation of dMSNs is sufficient in inducing a dip-rebound in ACh release as the stimulation pulses increase (Fig. 5J). Further analysis indicates that the dMSNs-induced dip-rebound reached the plateau at 0.5 of optical stimulation in vivo (Fig. 5K). The amplitude of dip and rebound in both dMSNs shows a positive correlation (Fig. 5L).

Next, we want to verify that the dMSN activation is required to induce the ACh dip-rebound in vivo. D1-Cre mice were infused with AAV-DIO-eNpHR plus AAV-gACh4m sensor as the experimental group and AAV-FLEX-tdT plus AAV-ACh4m as the control group, and optical fibers were implanted into the same sites. Two weeks after virus infusion, both groups underwent an FR1 schedule with a 100% chance of laser delivery along with reward delivery (Fig. 5M, 5N). After four weeks of training, both the control and experimental mice showed an immediate magazine entry after reward delivery (sFig. 6A, 6B), implying the establishment of action-outcome contingency and the motivation to collect rewards. However, analysis of active lever press and reward delivery learning curves showed that both groups acquired the task at similar rates, but the experimental group displayed a lower maximal level of active lever pressing and reward delivery compared with controls (sFig. 6C, 6D). The observation can be explained by the previous findings that dMSN activation is reinforcing(Kravitz, Tye et al. 2012, Cole, Robinson et al. 2018). Hence, optical inhibition of the dMSN dampened the reinforcing effect. Simultaneous fiber photometry recording of ACh dynamics on the last day of training revealed that optogenetic inhibition of dMSNs during reward delivery markedly attenuated the ACh dip–rebound pattern normally observed after lever presses (Fig. 5O-Q), confirming that post-action dMSN activation is required to induce the ACh dip-rebound in instrumental learning. Together, these results demonstrate that post-action dMSN activation is both required and sufficient for striatal ACh dip-rebound in instrumental learning.

**Figure 6.**
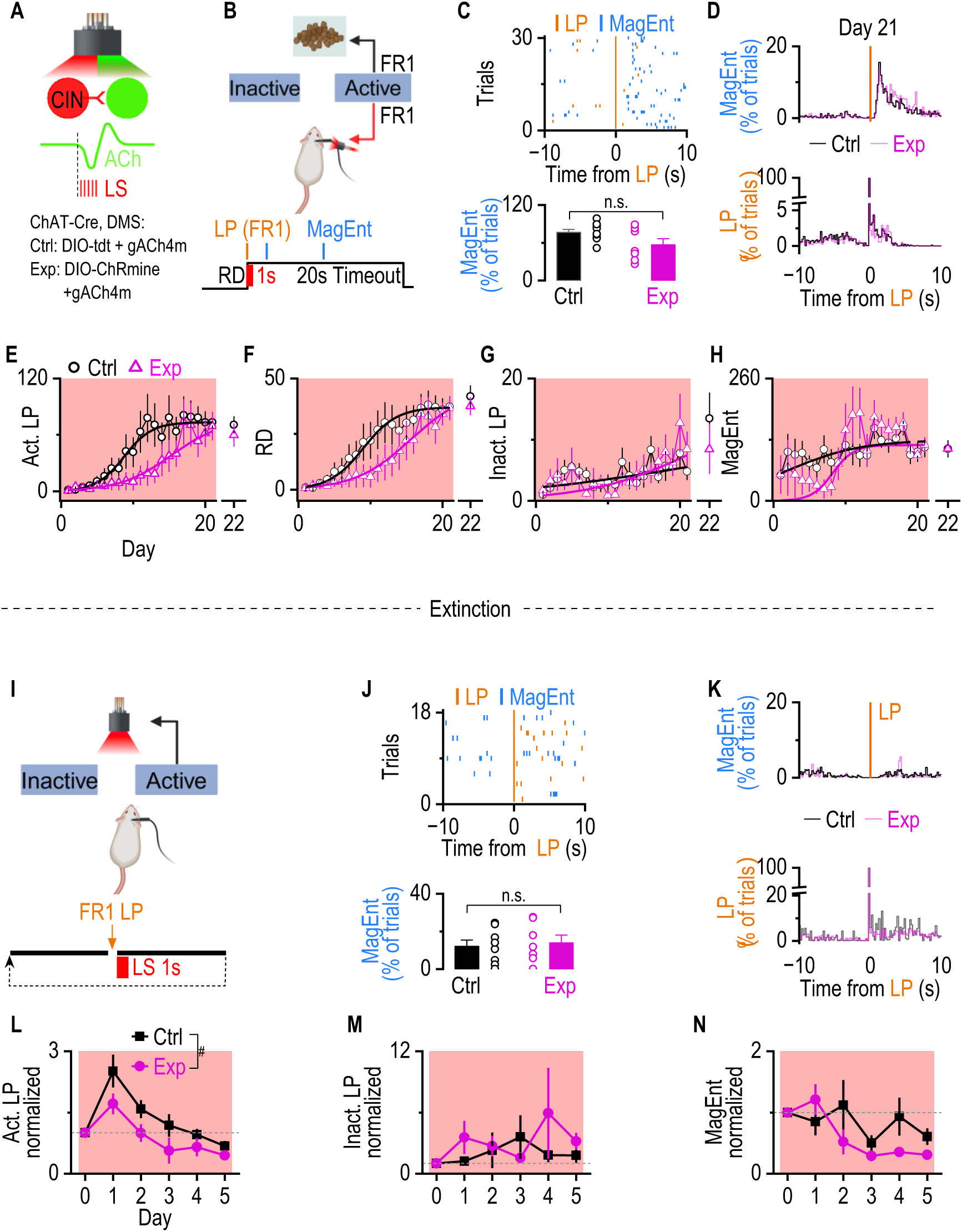
Optogenetic disruption of the ACh dip in the DMS impairs action-outcome learning and facilitates extinction. **A.** Experimental design: ChAT-Cre mice were infused with AAV-DIO-tdT plus AAV-gACh4m (control, Ctrl) or AAV-DIO-ChRmine plus AAV-gACh4m (experimental, Exp) into DMS. Immediately after each reward delivery, a 5-Hz light stimulation (LS, 615 nm, 1 sec) was delivered in both groups. **B.** Schematic of the behavioral, stimulating, and recording setup. Mice were trained in operant chambers under an FR1 schedule, where each active lever press (LP) triggered food delivery, 10 sec timeout, and light stimulation of CINs. **C.** Representative raster plots showing active lever presses and magazine entries aligned to reward delivery (Top). Immediate magazine entries after reward deliveries (within 3 sec) did not differ between the control and experimental groups (Bottom). n.s., not significant, *p* > 0.05 by unpaired t-test, n = 9 (Ctrl) and 7 (Exp) mice. **D.** Indistinguishable average distribution of Magazine entry (top) and lever presses(bottom) between Ctrl and Exp group. **E.** Active lever pressing (Act. LP) in the experimental group increased significantly more slowly over sessions than in the control group, indicating that ACh dip-rebound is required for the learning of A-O contingency. Act. LP across training days is shown for the Exp (magenta triangles) and Ctrl (black circles) groups. Solid curves represent the best-fit 3-parameter logistic regression (Ctrl group: K = 73.48 ± 1.87, x₀ = 8.71 ± 0.30, k = 0.521 ± 0.031, *R²* = 0.96; Exp group: K = 77.12 ± 9.55, x₀ = 15.35 ± 1.26, k = 0.278 ± 0.028, *R²* = 0.91). Wald *z*-tests indicated significant group differences in the inflection point (x₀) and slope (k) parameters, while asymptotic performance (*K*) did not differ (*zₓ₀* = 5.12, \*\*\**p* < 0.001; *zₖ = –5.82,* \*\*\**p* < 0.001*; z_K_ = 0.37, p* > 0.05). On the last day of training (Day 22), the laser will be removed. There is no difference in Act. LP between Day 22 and Day 21(Ctrl group: *t*_(6)_ = 1.23, *p* >0.05, paired t test; Exp group: *t*_(8)_ = 0.36, *p* >0.05, paired t test). **F.** Reward delivery (RD) in the Exp group increased significantly more slowly over sessions than that in the Ctrl group. RD across training days is shown for the Exp (magenta triangles) and Ctrl (black circles) groups. Solid curves represent the best-fit 3-parameter logistic regression (Ctrl group: K = 37.11 ± 1.11, x₀ = 8.96 ± 0.44, k = 0.43 ± 0.04, (*R²* = 0.97); Exp group: K = 45.41 ± 3.57, x₀ = 15.66 ± 0.91, k = 0.26 ± 0.02, (*R²* = 0.96)). Wald *z*-tests indicated significant group differences in the inflection point (x₀) and slope (k) parameters, while asymptotic performance (*K*) did not differ (*zₓ₀* = 2.95, \*\*\**p* < 0.001; *zₖ = –*2.16, \*\*\**p* < 0.001*; z_K_* = 0.97*, p* > 0.05). There is no difference in RD between Day 22 and Day 21(Ctrl group: *t*_(6)_ = 0.83, *p* >0.05, paired t test; Exp group: *t*_(8)_ = 1.19, *p* >0.05, paired t test). **G.** Inactive lever pressing (Inact. LP) in the Ctrl group and Exp group over sessions did not follow a clear sigmoidal trend (Ctrl group: K = 11.40 ± 39.03, x₀ = 22.00 ± 96.67, k = 0.07 ± 0.12, (*R²* = 0.23); Exp group: K = 16.01 ± 30.84, x₀ = 22.00 ± 24.85, k = 0.14 ± 0.09, (*R²* = 0.49)). **H.** The number of Magazine entries (MagEnt) in the Ctrl group and Exp group over sessions did not follow a clear sigmoidal trend (Ctrl group: K = 129.01 ± 12.99, x₀ = 2.00 ± 1.82, k = 0.20 ± 0.12, (*R²* = 0.28); Exp group: K = 118.39 ± 6.85, x₀ = 8.24 ± 0.99, k = 0.69 ± 0.48, (*R²* = 0.37)). **I.** Schematic of the behavioral, stimulating, and recording setup. Mice were trained in operant chambers under an FR1 schedule, the same as the schedule in panel A, except that there was no food delivered (pseudo reward delivery). **J.** Representative raster plots showing active lever presses and magazine entries aligned to pseudo reward delivery (Top). Immediate magazine entries after reward deliveries (within 3 sec) were diminished in the control and experimental groups (Bottom). n.s., not significant, *p* > 0.05 by unpaired t-test, n = 8 (Ctrl) and 7 (Exp) mice. **K.** Indistinguishable average distribution of Magazine entry (top) and lever presses(bottom) between Ctrl and Exp group aligned to pseudo reward delivery. **L-N.** Optical disruption of ACh dip-rebound facilitates extinction. **L.** Normalized active LP in the Exp group was significantly lower than in the Ctrl group across 5 days of extinction training (^#^*p* < 0.05, F(1, 14) = 4.97, by 2-way ANOVA with mixed effect analysis). **M.** Normalized inactive LPs between the Ctrl and Exp group were not different (*p* > 0.05, F(1, 14) = 0.50, by 2-way ANOVA with mixed effect analysis). **N.** Normalized magazine entries between the Ctrl and Exp group were not different (*p* > 0.05, F(1, 14) = 1.26, by 2-way ANOVA with mixed effect analysis).

### Disrupting the ACh dip-rebound slows down the establishment of contingency and facilitates extinction

To determine whether striatal ACh dip-rebound in the DMS is required for establishing the contingency, AAV-DIO-ChRmine plus AAV-ACh4m were infused into the DMS of ChAT-Cre mice as experimental groups or AAV-FLEX-tdT plus AAV-ACh4m as controls (Fig. 6A). We used the excitatory opsin ChRmine to activate CINs and offset the dip phase, which likely distorts the rebound and disrupts dip-rebound dynamics. Optical fibers were implanted at the same coordinates for simultaneous optogenetic stimulation and ACh recording. Two weeks after the infusion, both groups underwent the FR1 schedule with laser stimulation. Immediately after each reward delivery, both groups received optogenetic stimulation (5 Hz, 1 second, 615 nm) (Fig. 6B); this protocol was verified to disrupt the natural ACh dip-rebound, without inducing artificial bursting that could be aversive or cause motor disruption in a separated experimental group (sFig. 7, sFig. 8). After 21 sessions of FR1 instrumental training, both groups showed a stable immediate magazine entry rate after active lever presses, implying both control and experimental mice eventually learned the action-outcome contingency (Fig. 6C, 6D). However, logistic regression analysis showed a significant rightward shift and shallower slope of the active lever presses and reward delivery in the experimental group relative to controls (Fig. 6E, 6F). Removal of the light stimulation on Day 22 did not alter active LP or RD in either group (Fig. 6E, 6F). Inactive lever presses (Fig. 6G) and magazine entries (Fig. 6H) showed no consistent sigmoidal pattern (R² < 0.4). Those results suggest that disruption of the ACh dip-rebound slows down the establishment of A-O contingency.

We also assess the role of ACh dip-rebound in extinction learning, which requires the degradation of the original A-O contingency (Dickinson and Balleine 1994, Corbit and Balleine 2003, Bouton, Maren et al. 2021). If the ACh dip-rebound is required for maintaining the contingency, disrupting the dip-rebound should facilitate the extinction. To test this possibility, the same mice underwent 5 days of extinction training immediately after FR1 training, using the same laser stimulation protocol as in Fig. 6B to disrupt the ACh dip–rebound (Fig. 6I), except that no food rewards were delivered during extinction. During extinction training, both groups diminished the immediate magazine entries (Fig. 6J, 6K), consistent with previous findings that the disruption of ACh dip-rebound does not affect the reward-taking behavior. However, the number of lever presses in the experimental group was significantly lower than that in the control group (Fig. 6L). There was no difference in inactive lever presses and magazine entries between the experimental group and control group (Fig. 6M, 6N). Our results indicate that disrupting the ACh dip-rebound accelerates extinction, supporting the notion that ACh dip–rebound activity primarily serves to sustain the A-O contingency.

Together, our finding suggests that dMSNs-induced ACh dip-rebound is required for the establishment and maintenance of A-O contingency.

## Discussion

Previous circuit and slice-level studies have established a reciprocal relationship between dopaminergic terminals and CINs: optogenetic stimulation of DA terminals induces a CIN firing pause through D2 receptors(Chuhma, Mingote et al. 2014, Gallo, Greenwald et al. 2022), whereas CIN activation promotes DA release via nicotinic receptors on DA terminals(Surmeier and Graybiel 2012, Threlfell, Lalic et al. 2012, Liu, Cai et al. 2022). These findings formed a canonical view that DA inhibits ACh release, while ACh facilitates DA release. However, there is no solid in vivo data to support the findings so far(Taniguchi, Melani et al. 2024). Here, we found that in DMS, the ACh dip–rebound is driven by dMSN activity and primarily encodes the A–O contingency. In contrast, DA transients track reward outcomes, consistent with their role in encoding reward prediction errors(Schultz, Dayan et al. 1997, Waelti, Dickinson et al. 2001, Steinberg, Keiflin et al. 2013, Amo 2024). Our results also indicate that ACh and DA do not exert obligatory modulation on each other in every behavioral context. Their signals diverge whenever the functional demands of the behavior differ. For example, during reward delayed tasks (Fig. 3), ACh and dMSN signals shift in time while DA amplitude and timing remain stable. Conversely, in unrewarded trials, the ACh rebound is significantly larger than in rewarded trials, potentially reflecting local DA disinhibition on ACh rebound. Together, these observations suggest that DA–ACh interactions are conditional rather than universal, emerging only in contexts where both neuromodulators are co-engaged by behavioral structure.

In Fig. 3, the reward delivery was temporally separated from level presses, and reward delivery itself triggered dMSNs activity and the ACh dip-rebound after 6 sessions of training. A similar phenomenon was also observed in the contingency degradation schedule. A potential explanation is that reward delivery may elicit additional behavior related to the potential outcome, but was not detected by our behavior chamber system. In this case, this dMSN activity and the ACh dip-rebound were still tracking the A-O contingency. Another possibility is that despite efforts to minimize Pavlovian contributions, the mechanical cue associated with pellet delivery may have acquired predictive value. Thus, the dMSN activity and ACh dip–rebound observed at reward delivery may reflect not only the A–O contingency but also cue–outcome contingencies formed incidentally during training. This hypothesis is also supported by other studies showing that visual stimuli are sufficient to induce striatal dMSN activation(Cui 2013, Hannah C. Goldbach1-4 2025).

A consistent feature across multiple behavioral schedules was a prolonged suppression of both dMSN and iMSN activity in reward delivery trials. This inhibition cannot be explained by DA release, as D_1_ receptors in dMSNs are excitatory, and the suppression was observed in both dMSNs and iMSNs. Instead, several alternative circuit mechanisms may contribute: 1) local GABAergic interneurons, such as Lhx6+ or fast-spiking interneurons, which strongly inhibit MSNs(Tepper, Koos et al. 2004, O’Hare, Ade et al. 2016); 2) Inhibitory feedback from Globus Pallidus Externa arkypallidal neurons(Mallet, Schmidt et al. 2016, Baker, Kang et al. 2023). Its functional relevance remains an open question.

## Methods

### Viral injection and implantation of fiber optic ferrules

Anesthesia was induced with 0.5 – 1.0% isoflurane through a nose cone mounted on a stereotaxic apparatus (Kopf Instruments). The scalp was opened, and bilateral holes were drilled in the skull (+0.3mm AP, ±1.65mm ML from Bregma). Virus was injected into the left and right dorsomedial striatum −3.0mm DV from the top of the brain through a 33-gauge steel injector cannula (Plastics1) using a syringe pump (World Precision Instruments) over 10 min. The injection cannula was left in place for 10 min following the injection, and then slowly removed. After the viral injection, two plastic mounts containing two fibers (105µm core/125µm cladding) mounted in 1.25mm zirconia ferrules were slowly lowered into the brain and cemented in place such that each fiber was aimed at the dorsomedial striatum on either side. To allow time for viral expression, animals were housed for at least 2 weeks following injections before any experiments were initiated. All surgical procedures were performed under aseptic conditions.

| Virus | Vendor | Lot# |
| --- | --- | --- |
| pGP-AAV-syn-FLEX-jGCaMP7f-WPRE | Addgene | 104492-AAV9 |
| rAAV5/PAAV-Ef1a-DIO-hsChRmine-Oscarlet | UNC | AV9622 |
| rAAV2/9-hsyn-DA3h | BrainVTA | PT-4721 |
| rAAV2/9-hSyn-gACh4m | BrainVTA | PT-7021 |
| rAAV5/Ef1a-DIO-eNPHR3.0-mCherry-WPRE | UNC | AV4832G |

### Fiber photometry

The fiber photometry experiment was performed using a bundle-imaging fiber photometry setup (Neurophotometrics FP3002 V2). Genetically encoded sensors were stimulated using 415-nm (isosbestic) and 470-nm excitation LED. The LED power coming out of the ferrule is between 50 and 100 μW. Before the recording, mice were tethered to a fiber optic cable connected to the implanted optic fiber for the later fiber photometry recording and placed in the behavior chamber for the behavior task. Two minutes after the fiber photometry recording system was turned on with a 25 Hz sample rate and the LEDs were delivered, the behavior task was initiated. At the beginning of the behavior task, a TTL signal will be delivered from the behavior task system to the fiber photometry recording system. The TTL signal was used to align the behavior data with the fiber photometry data. After the behavior task, the fiber photometry recording will be turned off, and the mice will be returned to their home cage. All the behavior data and fiber photometry data will be saved for further analysis with MATLAB scripts. All datasets and custom MATLAB scripts will be available upon reasonable request. Scripts are available via GitHub (https://github.com/Chen-RuiFeng/Fiber-Photometry_NeuroPhotometrics).

### Electrophysiology

Coronal slices (250 µm) containing the dorsomedial striatum were cut at 0.14 mm/s in ice-cold cutting solution (in mM: 40 NaCl, 148.5 sucrose, 4.5 KCl, 1.25 NaH₂PO₄, 25 NaHCO₃, 0.5 CaCl₂, 7 MgSO₄, 10 dextrose, 1 sodium ascorbate, 3 myo-inositol, 3 sodium pyruvate) saturated with 95% O₂/5% CO₂. Slices recovered for 45 min in a 1:1 mixture of cutting and external solution (in mM: 125 NaCl, 4.5 KCl, 2.5 CaCl₂, 1.3 MgSO₄, 1.25 NaH₂PO₄, 25 NaHCO₃, 15 sucrose, 15 glucose). For recordings, the bath was maintained at 32 °C (2–3 mL/min perfusion). Potassium-based internal solution (in mM: 123 K-gluconate, 10 HEPES, 0.2 EGTA, 8 NaCl, 2 MgATP, 0.3 NaGTP; pH 7.3 with KOH) was used for current-clamp. Glutamatergic transmission was blocked with NBQX (10 µM) + AP5 (50 µM), and GABAergic transmission was blocked with picrotoxin (100 µM). CINs were identified by fluorescence, size, spontaneous firing, and membrane sag (held at –60 mV); dMSNs were held at –70 mV. Data were acquired with Clampex (pClamp 10.7, Molecular Devices) and analyzed using Clampfit and Mini Analysis (Synaptosoft).

### Behavioral Assays

All behavior tests in the work were combined with fiber photometry recording. Animals will experience one 30-minute behavior session on every workday. The behavior data and fiber photometry data will be collected at the same time and will be analyzed with MATLAB scripts. After daily behavior training, food will be applied to maintain the body weight.

### FR1 schedule

Mice were placed in operant chambers (MED Associates Inc.) equipped with two levers (left and right levers) and a food delivery port. During training sessions, the levers remained continuously available to prevent lever withdrawal from serving as a potential reward-related cue. Pressing the active lever (right lever) resulted in the immediate delivery of a food pellet as a reinforcer. Following each active lever press, a 20-second timeout period was imposed, during which additional lever presses did not result in reward delivery. The left lever remained inactive throughout the session, serving as a control (sFig. 2A).

### VR2 schedule

A Variable Ratio 2 (VR2) schedule was introduced following several weeks of FR1 training. Mice were placed in the same operant chambers as before, with the right lever designated as the active lever and the left lever serving as the inactive lever. Under the VR2 schedule, each active lever press had a 50% probability of triggering food pellet delivery and a 50% probability of no reward. Hence, within each session, lever presses were categorized into rewarded trials (presses that resulted in food pellet delivery) and unrewarded trials (presses that did not yield a reward). The rewarded trials and unrewarded trials are distributed in a random order. Regardless of whether a reward was delivered, a 20-second timeout period followed each active lever press. During the timeout period, two levers are withdrawn to prevent continuous responding (sFig. 2B).

### FR1-FD5 schedule

With the same cohort of mice that experienced VR2 training, mice underwent 3 days of FR1 training before transitioning to the Fixed Ratio 1 - Fixed Delay 5 seconds (FR1-FD5) schedule. Mice were placed in the same operant chambers as before, with the right lever designated as the active lever. Under this schedule, each active lever press initiated a 5-second fixed delay before the guaranteed delivery of a food pellet (100% reward probability). This delay was implemented to temporarily separate the lever press (LP) from reward delivery (RD). Following each active lever press, a 20-second timeout period was imposed, which included the 5-second delay. During this timeout, both levers were withdrawn to prevent continuous responding and ensure distinct trial separations (sFig. 2C).

### Contingency degradation schedule

With the same cohort of mice that experienced FR1-FD5 training, mice underwent 3 days of FR1 training before transitioning to the contingency degradation schedule. Mice were placed in the same operant chambers as before. Under the contingency degradation schedule, lever presses no longer resulted in reward delivery. Instead, each active lever press triggered a 20-second timeout period, during which no reinforcer was available. Food pellets were delivered randomly, independent of lever presses, with an average inter-reward interval of approximately 80 seconds. The 80-sec interval was selected because previous data indicated a lower quartile of 22 rewards delivered within 30 min at this rate (30*60/22 ≍ 81.82). This schedule was designed to disrupt the action-outcome contingency (sFig. 2D).

### Food deprivation

Food deprivation is applied to all animals that are experiencing the series of training (sFig 2e). A fixed daily food (2 g/mouse/day) was provided to maintain 85% of baseline weight. If the mice did not learn to finish the task to achieve food pellets for 3 consecutive days, 1 g/mouse/day extra food will be added. The body weight was measured, and behavioral performance was checked daily.

### Three-parameter logistic regression

To quantify group learning curves, behavioral performance across training days was fitted with a three-parameter logistic function of the form:

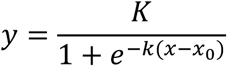

where *y* represents the behavioral performance (e.g., lever presses, reward delivery, inactive lever presses, or magazine entry per session), and *x* denotes the training day. The parameters *K*, *x₀*, and *k* correspond to the asymptotic performance level, the inflection point (the day at which performance reaches half of the asymptote), and the slope (learning rate) of the curve, respectively. The lower asymptote was fixed at zero. For each experimental group, the daily behavioral data were fitted using nonlinear least-squares optimization with MATLAB scripts. For group comparisons, parameter estimates (*K*, *x₀*, *k*) obtained from each group were statistically compared using two-sample Wald z-tests based on their estimated standard errors, with significance defined as *p* < 0.05.

### Quantification of immediate magazine entries

For every rewarded trial, if a magazine entry occurred within 3 seconds after the onset of reward delivery, it was counted as an immediate magazine entry( 𝑁_immediate entries_). The total number of immediate entries was then divided by the total number of rewarded trials (𝑁_rewarded trials_), yielding the percentage of trials with immediate magazine entry:

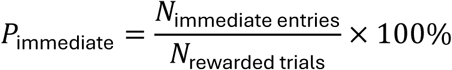

## Acknowledgments

This research was supported by NIH grants R01AA027768 (J.W.) and U01AA025932 (J.W.).

## Author contributions

J.W., R.C., and H.G. initiated the project. X.X. and R.C. designed the behavior experiments. R.C. performed the behavioral experiments, fiber photometry experiments, and optogenetic stimulations with fiber photometry and analyzed the corresponding data. H.G. performed the electrophysiological experiments and analyzed the corresponding data. R.C. performed the histology experiments. X.W. performed animal breeding for all experiments. J.W. and R.C. wrote the manuscript with substantial input from X.X.

## Supplementary

**Supplementary Figure 1.**
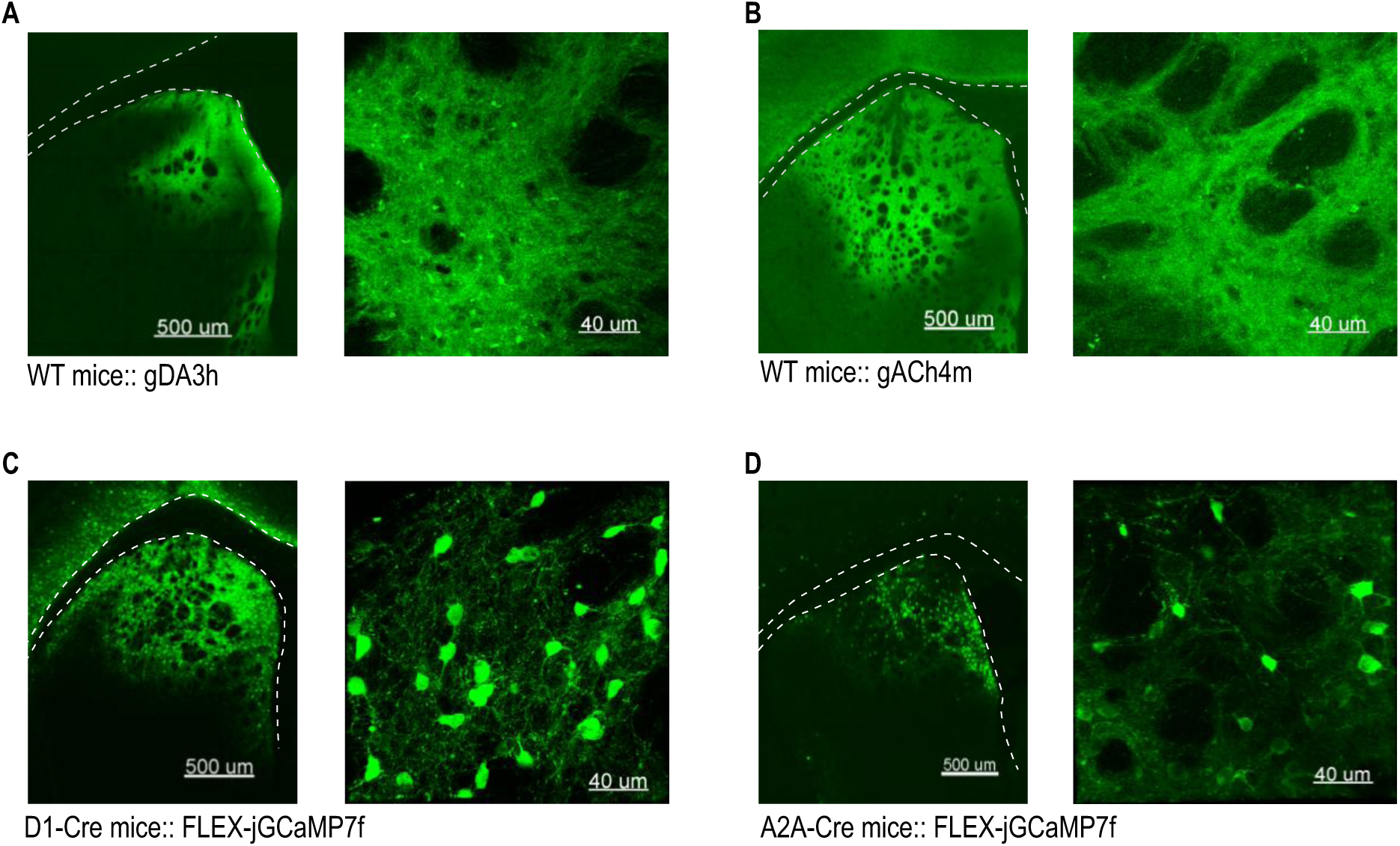
Histology for gACh4m, gDA3h, and jGCaMP7f expression in D1-Cre mice or A2A-Cre mice in the DMS. **A.** Confocal images of gACh4m expression for a representative mouse recorded for Fig.1. The Left image shows the expression of gACh in the DMS. The right image shows the enlarged detail of gACh expression. **B.** Confocal images of gDA3h expression for a representative mouse recorded for Fig.1. The Left image shows the expression of gDA3h in the DMS. The right image shows the enlarged detail of gDA3h expression. **C.** Confocal images of Cre-dependent jGCaMP7f expression for a representative D1-Cre mouse recorded for Fig.1. The Left image shows the expression of jGCaMP7f in the DMS. The right image shows the enlarged detail of jGCaMP7f expression. **D.** Confocal images of Cre-dependent jGCaMP7f expression for a representative A2A-Cre mouse recorded for Fig.1. The Left image shows the expression of jGCaMP7f in the DMS. The right image shows the enlarged detail of jGCaMP7f expression.

**Supplementary Figure 2.**
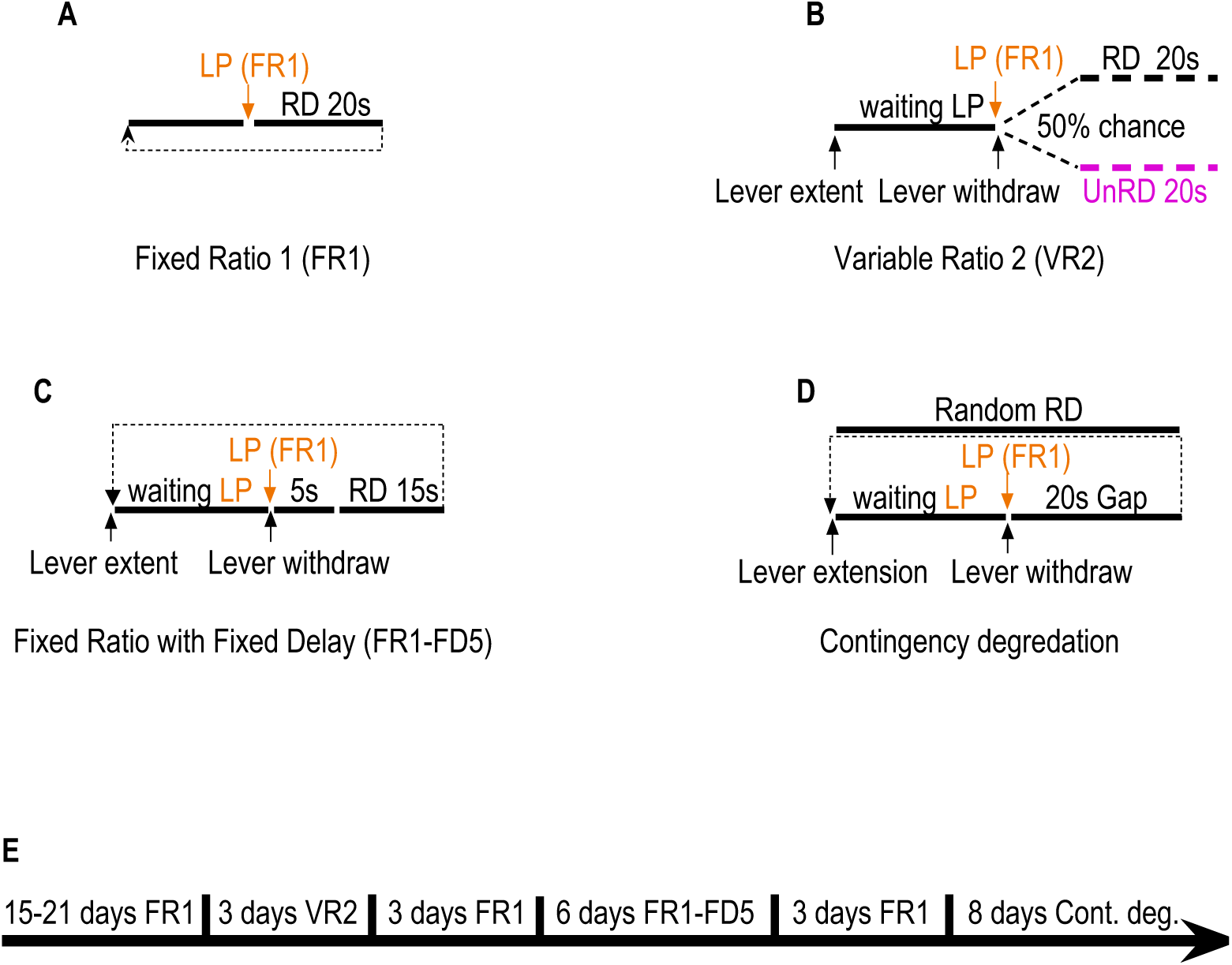
Behavioral task schedules and training timeline. (**A**–**D**) Schematic diagrams of the four operant conditioning schedules used across training phases. More details can be found in the Method section **A. Fixed Ratio 1 (FR1):** A single lever press (LP) after lever extension (L extent) triggered immediate reward delivery (RD; 20 s timeout). **B. Variable Ratio 2 (VR2):** Each lever press had a 50% probability of resulting in reward delivery (RD; 10 s timeout) or no reward (non-RD; 10 s timeout). **C. Fixed Ratio with Fixed Delay (FR1–FD5):** A single lever press initiated a fixed 5-second delay, after which the reward was delivered (RD; 10 s timeout). **D. Fixed Ratio with Variable Interval (FR1–VI80):** lever presses initiated a 10-second timeout. Reward was delivered with a variable time interval (∼80 s). **E.** Timeline of the behavioral training protocol. Animals were initially trained under the FR1 schedule for 15–21 days, followed sequentially by VR2 (5 days), FR1–FD5 (6 days), a brief reversion to FR1 (4 days), and finally FR1–VI80 (8 days).

**Supplementary Figure 3.**
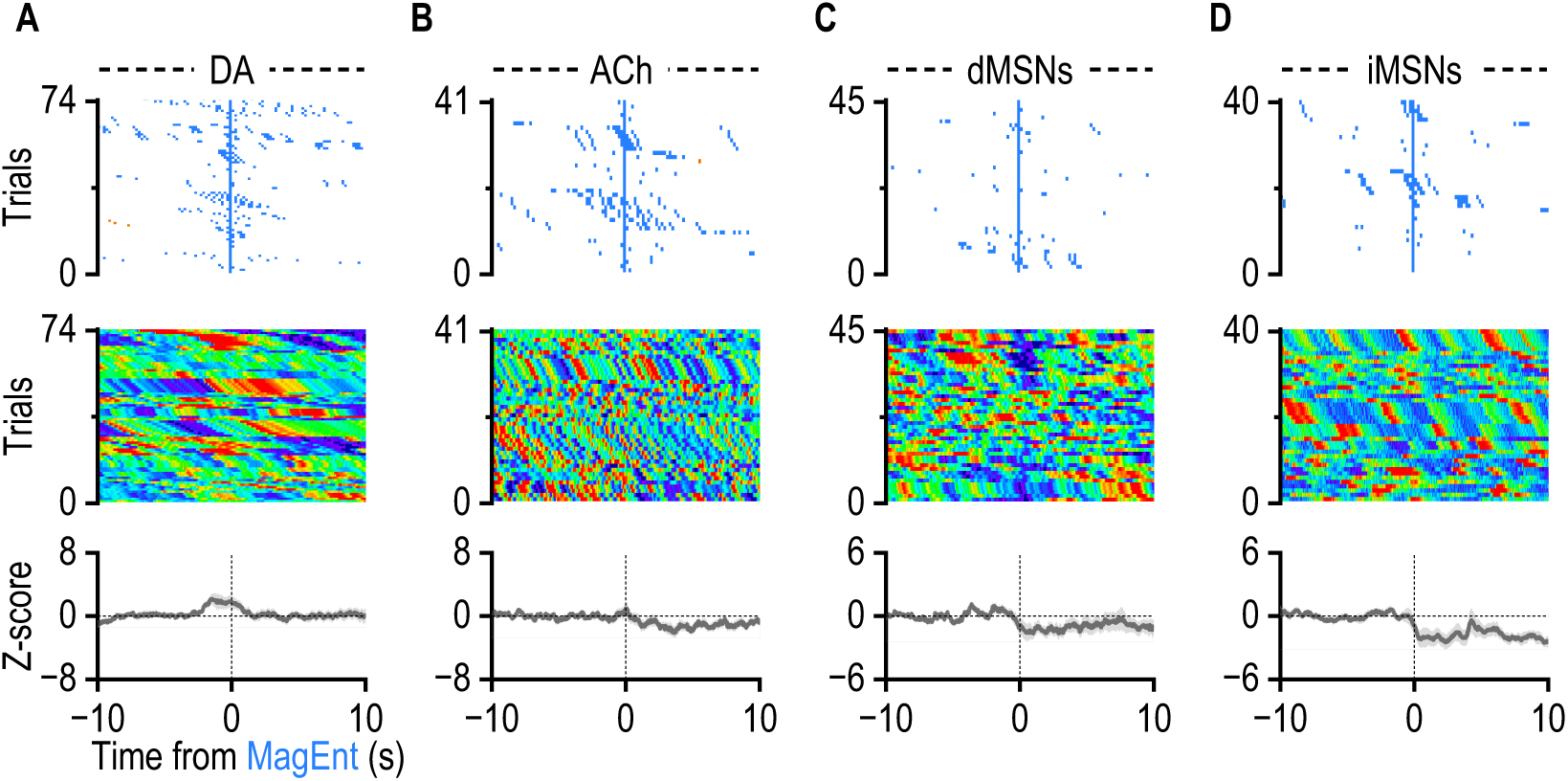
ACh, DA, dMSN, and iMSN have no significant response to magazine entry in naïve mice. **A-D**. (Top) Sample Raster plots showing the distribution of lever presses and magazine entries in naïve mice from 4 groups(1, wild-type mice infused with AAV-gACh; 2, wild-type mice infused with AAV-gDA; 3, D1-Cre mice infused with AAV-Flex-jGCaMP7f; 4, A2A-Cre mice infused with AAV-Flex-jGCaMP7f) aligned to magazine entries. (Middle) Sample heatmap showing neuronal activity or neurotransmitter dynamics in the same session as Top, aligned to the magazine entries. (Bottom) average neuronal activity or neurotransmitter dynamics of four groups aligned to magazine entries.

**Supplementary Figure 4.**
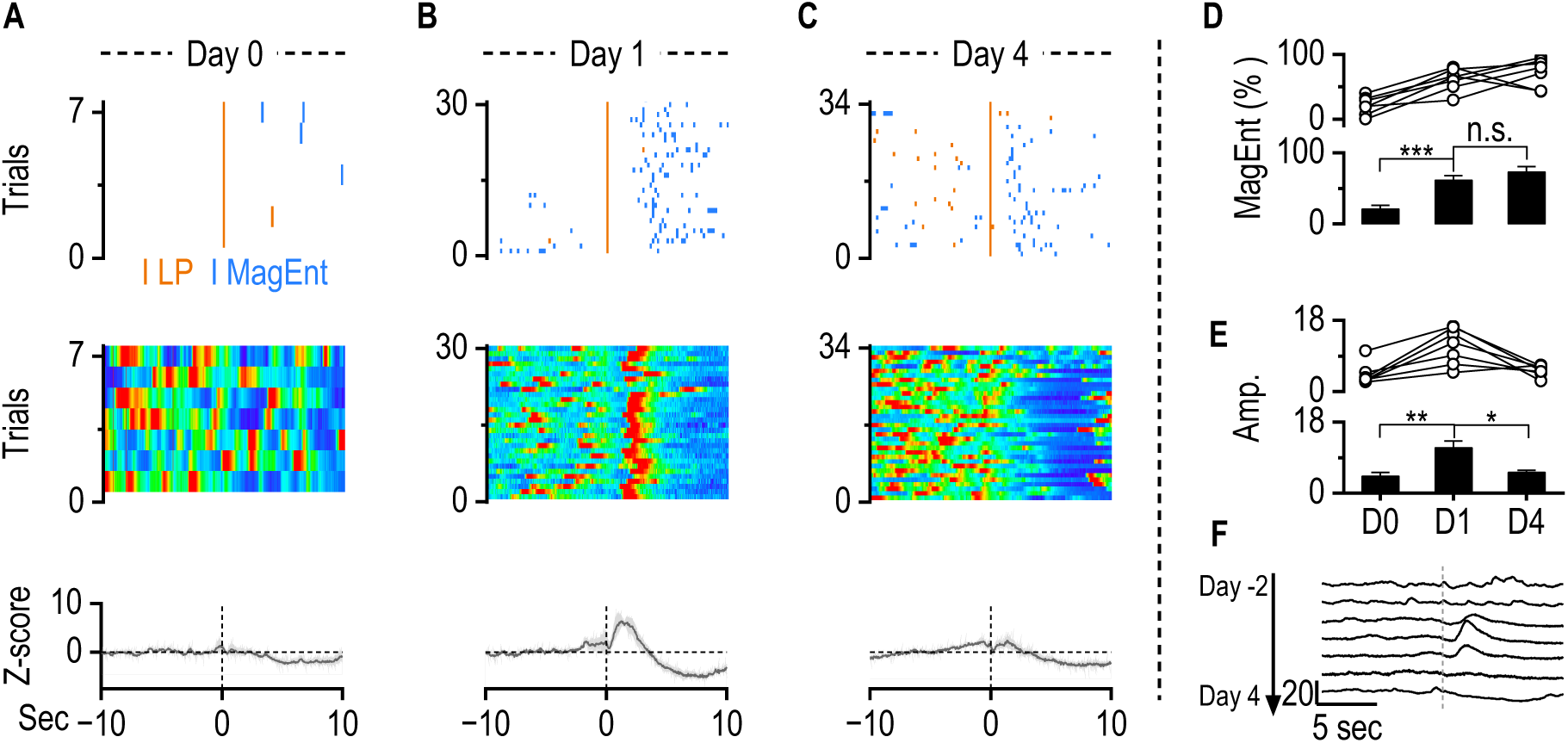
iMSN activity emerges during initial contingency learning and diminishes with continued training. At the beginning of FR1 training, before the animals had learned the action-outcome contingency, lever presses did not evoke any detectable iMSN activity. However, after several days of training, during the initial phase when mice first learned the contingency, iMSNs showed increased activity both before and after lever presses. With continued training over the following four days, this iMSN activity gradually diminished. (A–C) Representative behavioral and photometry data from a sample A2A-Cre mouse expressing jGCaMP7f in the DMS and trained under an FR1 schedule. Top: raster plots showing lever presses (orange) and magazine entries (blue) across trials. Middle: trial-aligned heatmaps of iMSN calcium signals time-locked to lever press (time = 0). Bottom: population average iMSN activity (Z-scored ΔF/F) across 7 mice. **A.** Day 0: prior to learning, lever presses were not consistently followed by immediate magazine entries, and iMSNs showed no activity change. **B.** Day 1: the first day the mouse exhibited learned contingency between lever press and reward; magazine entries reliably followed lever presses, and iMSNs showed increased activity around lever presses. **C.** Day 4: While immediate magazine entries persisted, iMSN activity following lever presses was reduced. **D.** Quantification of the percentage of trials with immediate magazine entries (within 2 s after lever press) shows significant increases on Day 1 and Day 4 compared to Day 0 (n = 7 mice; Day 0 vs Day 1: *t*_(6)_ = 6.09, ***p < 0.001; Day 1 vs Day 4: *t*_(6)_ = 1.00, p > 0.05, paired t test). **E.** Amplitude of iMSN activity following lever press was significantly elevated on Day 1 compared to Day 0 and Day 4 (Day 0 vs Day 1: *t*_(6)_ = 4.71, **p < 0.01; Day 1 vs Day 4: *t*_(6)_ = 3.22, *p < 0.05, paired t test). **F.** Example continuous trace of iMSN activity from one mouse across multiple training days (Day -2 to Day 4), showing transient activation emerging on the day of learning and declining thereafter.

**Supplementary Figure 5.**
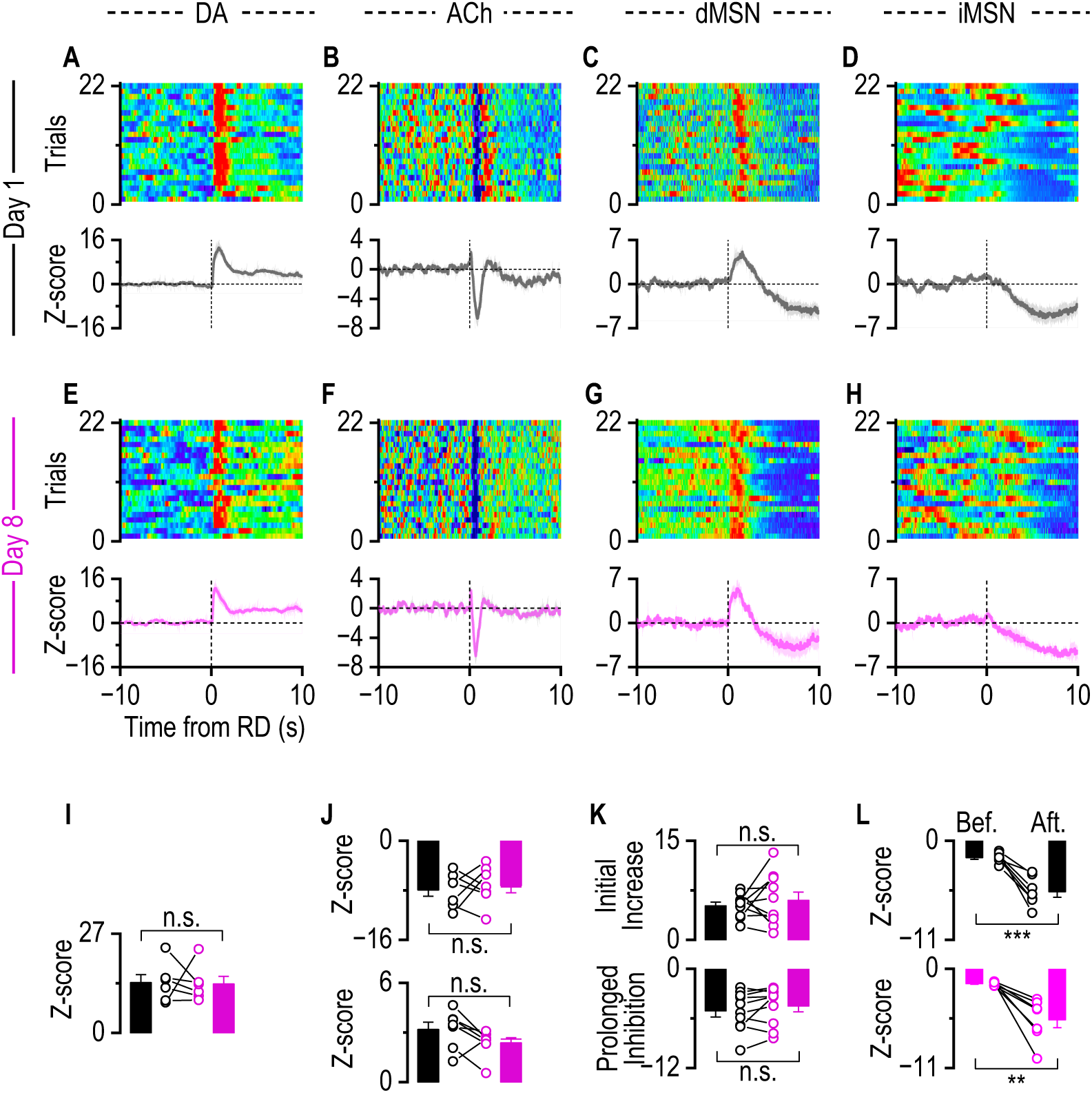
Contingency degradation does not change striatal ACh, DA, dMSN, and iMSN response to reward delivery. Mice underwent an 8-day contingency degradation protocol in which lever presses no longer led to reward delivery; instead, rewards were delivered non-contingently at ∼80-second variable intervals. (A–D) Neural dynamics on Day 1 of contingency degradation training. **A.** Top: Heatmap of ACh signals across trials aligned to reward delivery (time 0) showing a brief burst followed by a dip-rebound pattern. Bottom: Average ACh response. **B.** Top: Heatmap of DA signals showing a robust increase following reward delivery. Bottom: Average DA response. **C.** Top: Heatmap of dMSN calcium activity showing an increase followed by a sustained decrease after reward delivery. Bottom: Average dMSN response. **D.** Top: Heatmap of iMSN calcium activity showing a reduction following reward delivery. Bottom: Average iMSN response. **E–H.** Same as panel **A–D**, but for Day 8 of contingency degradation training. DA, ACh, dMSN, and iMSN responses remained qualitatively similar to Day 1, indicating persistent neural responses to reward delivery. **I–L.** Quantification of signal amplitudes for DA increase (**I**), ACh dip and rebound (**J**), and MSN activity (**K**, **L**) on Day 1 (black) and Day 8 (magenta), showing no significant difference across days (DA: *t*_(5)_ = 0.10, *p* >0.05, paired t test; ACh dip: *t*_(6)_ = 0.47, *p* >0.05, paired t test; ACh rebound: *t*_(6)_ = 1.98, *p* >0.05, paired t test; dMSN initial activity increase: *t*_(9)_ = 0.80, *p* >0.05, paired t test; dMSN prolonged inhibition: *t*_(9)_ = 1.41, *p* >0.05, paired t test; iMSN: Day 1 before RD vs. after RD, *t*_(6)_ = 6.50, \*\*\**p* <0.001, paired t test; Day 8 before RD vs. after RD, *t*_(6)_ = 4.47, \*\**p* <0.01, paired t test;).

**Supplementary Figure 6.**
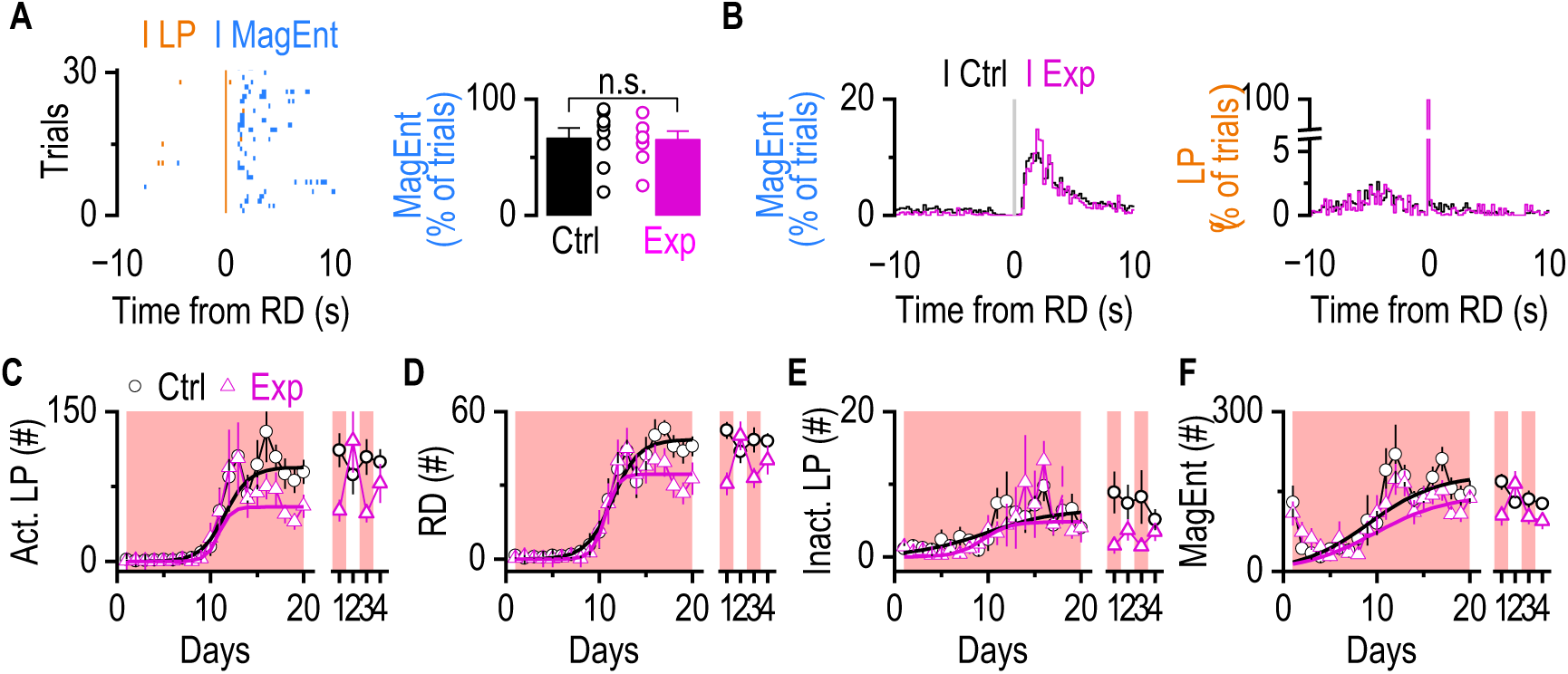
dMSN inhibition after lever press decreased the reinforcing effect. To test the functional role of dMSN activity after reward delivery in instrumental learning, and to verify that the optical inhibition of dMSN is sufficient to diminish the ACh dip-rebound. D1-Cre mice infused with DIO-ChRmine and gACh sensor underwent an FR1 schedule with a 100% chance of laser delivery after reward delivery. **A.** Representative raster plots showing active lever presses and magazine entries aligned to reward delivery (Left). Immediate magazine entries after reward deliveries (within 3 seconds) did not differ between the control and experimental groups (Right). n.s., not significant, p > 0.05 by unpaired t-test, n = 8 (Ctrl) and 8 (Exp) mice. **B.** Average distribution of Magazine entry (top) and lever presses(bottom) between Ctrl (black) and Exp (red) group. **C.** The Exp group showed higher maximal Active lever pressing (Act. LP) than the Ctrl group, despite similar acquisition dynamics. Solid curves represent the best-fit 3-parameter logistic regression (Ctrl group: K = 94.10 ± 5.76, x₀ = 11.57 ± 0.59, k = 0.77 ± 0.12, (R² = 0.89); Exp group: K = 54.40 ± 4.39, x₀ = 10.87 ± 0.81, k = 1.50 ± 0.64, (R² = 0.73)). Wald z-tests indicated significant group differences in asymptotic performance (K), while the inflection point (x₀) and slope (k) did not differ (zₓ₀ = -0.70, *p* > 0.05; zₖ = 1.13, *p* > 0.05; z_K_ = -5.48, \*\*\**p* < 0.001). On Day 22 and Day 24, the laser will be removed. There is no difference in Act. LP between Laser off sessions and Laser on sessions in the Ctrl group, but in the experimental group, removing the laser stimulation significantly increased Act. LP (Ctrl group: *t*_(7)_ = 0.25, *p* >0.05, paired t test; Exp group: *t*_(6)_ = 2.6, \**p* <0.05, paired t test). **D.** The Exp group showed higher maximal reward delivery (RD) than the Ctrl group despite similar acquisition dynamics (Ctrl group: K = 48.59 ± 1.89, x₀ = 11.43 ± 0.46, k = 0.71 ± 0.09, (R² = 0.96); Exp group: K = 34.48 ± 1.81, x₀ = 10.62 ± 0.39, k = 1.50 ± 0.37, (R² = 0.94)). Wald z-tests indicated significant group differences in asymptotic performance (K), while the inflection point (x₀) and slope (k) did not differ (*z*ₓ₀ = -1.35, *p* > 0.05; *z*ₖ = 2.10, *p* > 0.05; *z*_K_ = -5.39, \*\*\**p* < 0.001). Same as Act. LP, there is no difference in RD between Laser off sessions and Laser on sessions in the Ctrl group, but in the experimental group, removing the laser stimulation significantly increased Act. LP (Ctrl group: *t*_(7)_ = 0.09, *p* >0.05, paired t test; Exp group: *t*_(6)_ = 2.78, \**p* <0.05, paired t test). **E.** Inactive lever pressing (Inact. LP) in the Ctrl group and Exp group over sessions did not follow a clear sigmoidal trend (Ctrl group: K = 6.57 ± 1.64, x₀ = 9.82 ± 2.95, k = 0.26 ± 0.13, (R² = 0.56); Exp group: K = 6.63 ± 4.53, x₀ = 13.28 ± 8.37, k = 0.23 ± 0.17, (R² = 0.33)). **F.** The number of Magazine entries (MagEnt) in the Ctrl group and Exp group over sessions did not follow a clear sigmoidal trend (Ctrl group: K = 180.94 ± 28.59, x₀ = 9.21 ± 1.95, k = 0.27 ± 0.08, (R² = 0.54); Exp group: K = 143.20 ± 28.79, x₀ = 9.79 ± 2.72, k = 0.26 ± 0.14, (R² = 0.18)).

**Supplementary Figure 7.**
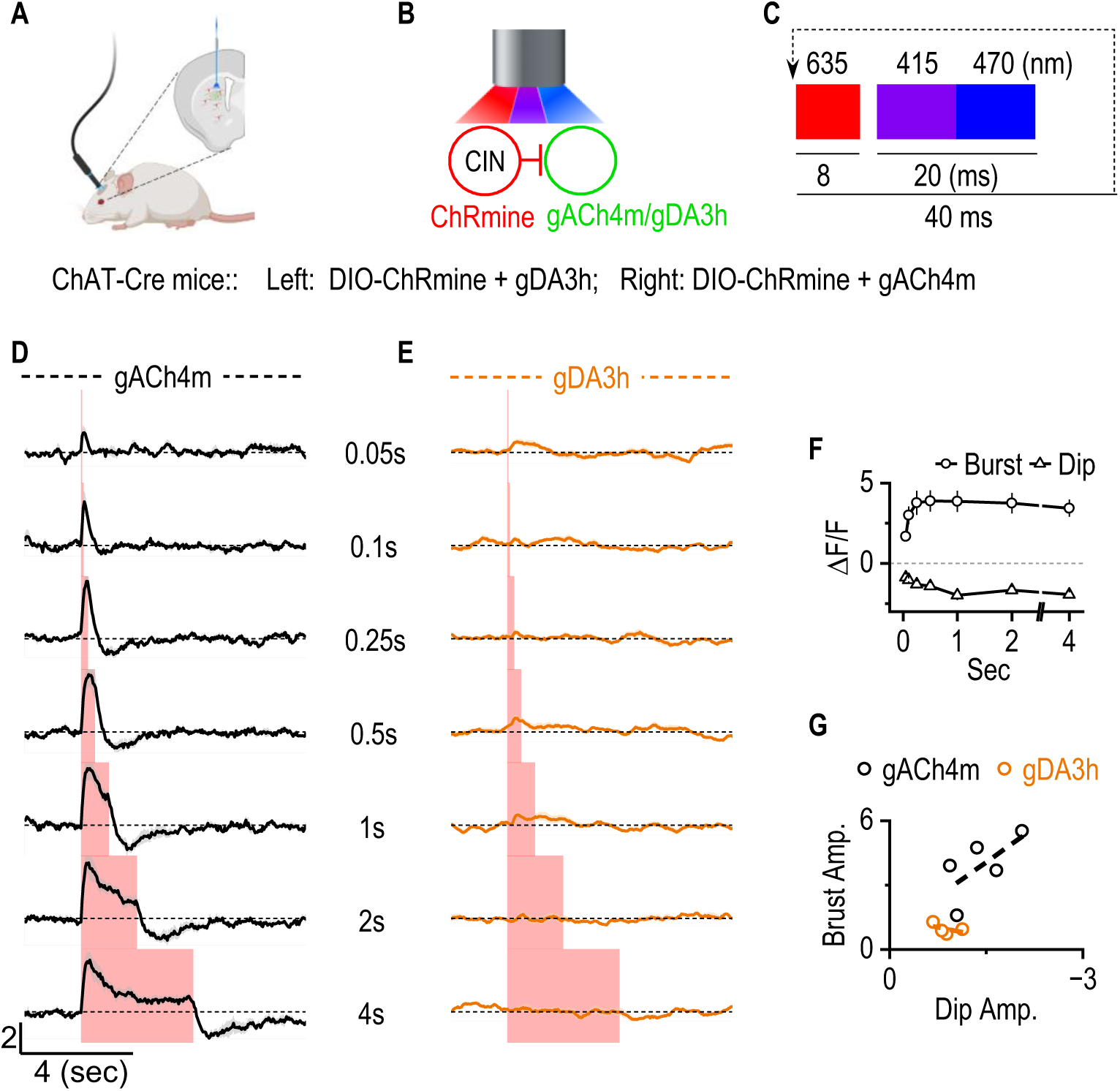
Optical stimulation of CIN induced an ACh burst-dip in vivo and no response in DA release. **A.** Diagram of the combined experimental setup for ChAT-Cre mice, showing simultaneous optogenetic stimulation and fiber photometry to assess DA release, as a control, or ACh release, as the experimental group, following CIN activation. In the control group, ChAT-Cre mice were infused with AAV-DIO ChRmine and AAV-gACh4m. In the Experimental group, ChAT-Cre mice were infused with AAV-DIO-ChRmine and AAV-gDA3h. **B.** Simplified circuit diagram depicting Cholinergic input from CIN onto other neurons expressing GRAB_gACh4m_ or GRAB_gDA3h_. **C.** Stimulation protocol: A cyclic sequence of three excitation wavelengths was used—635 nm (8 ms) for optogenetic activation of ChRmine triggered by TTL input, 415 nm (10 ms) as a photometry reference, and 470 nm (10 ms) for excitation of gACh4m. The full cycle lasted 40 ms and was continuously repeated to enable simultaneous optogenetic stimulation and real-time monitoring of ACh dynamics via fiber photometry. **D.** In vivo fiber photometry traces of ACh dynamics during optogenetic stimulation of CIN at increasing durations (0.05, 0.1, 0.25, 0.5, 1, 2, and 4s). ACh burst and dip become more pronounced with longer CIN stimulation (n = 5 mice). **E.** In vivo fiber photometry traces of DA dynamics during optogenetic stimulation of CIN at increasing durations (0.05, 0.1, 0.25, 0.5, 1, 2, and 4s). DA has no response to CIN activation, implying that even though DA terminals express nicotinic receptors, ACh dynamics did not affect DA release in vivo (n = 4 mice). **F.** Quantification of ACh Burst and dip amplitudes evoked by optogenetic stimulation of CIN at varying durations. The burst and dip amplitudes increased with stimulation duration and peaked at 0.5 seconds. **G.** A negative correlation between ACh burst and dip amplitudes was observed following CIN stimulation (*r* = -0.66, R^2^ = 0.44, slope = -2.13). Each dot represents one mouse in response to 25Hz, 0.5-second stimulation.

**Supplementary Figure 8.**
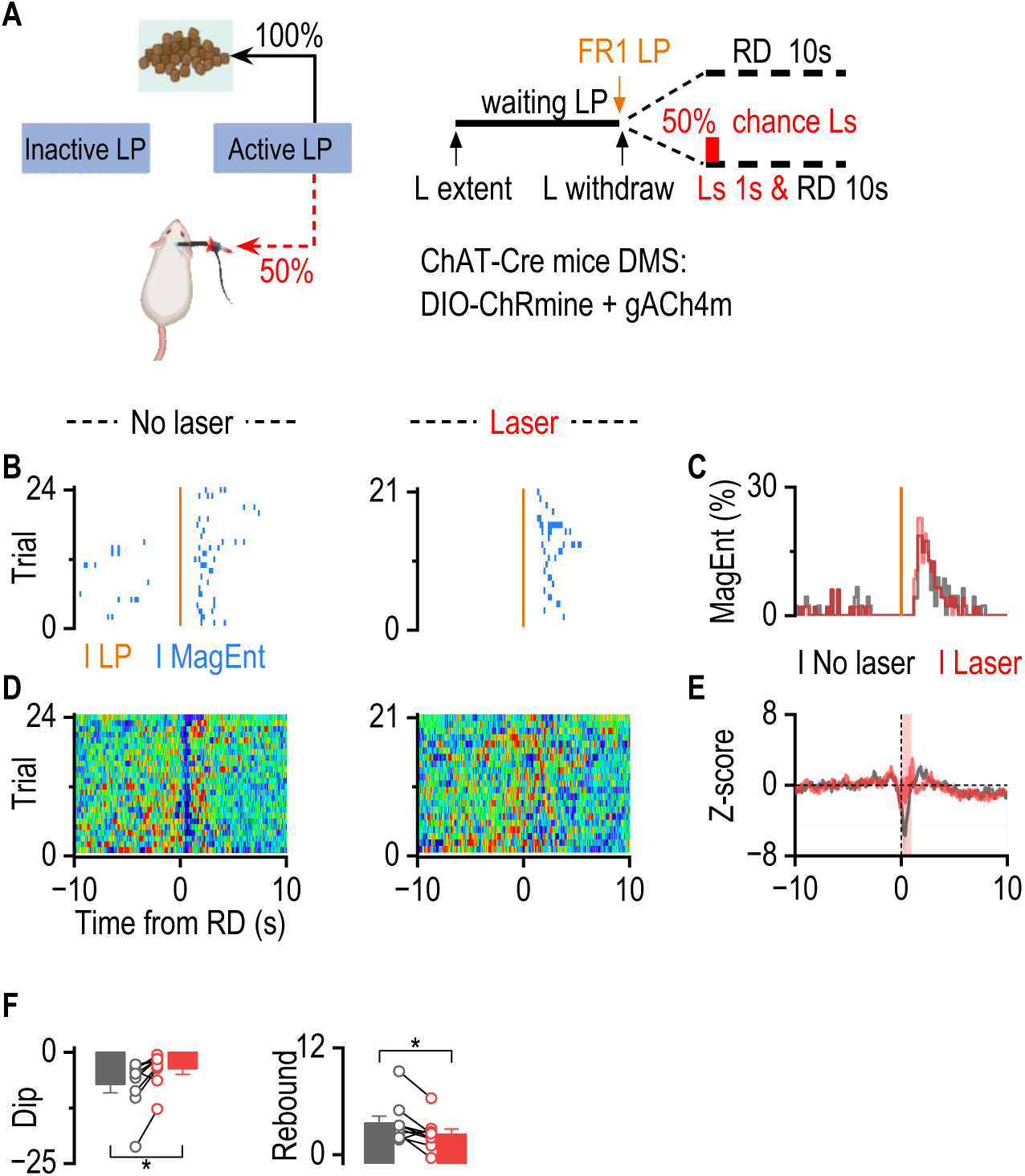
Optical stimulation of CIN after reward delivery is sufficient to disrupt ACh dip-rebound. To verify that the optical stimulation of CIN is sufficient to diminish the ACh dip-rebound. Well-trained ChAT-Cre mice infused with DIO-ChRmine and gACh4m und a FR1 schedule with a 50% chance of laser delivery after reward delivery. **A.** Schematic of the behavioral paradigm. Mice were tested under an FR1 schedule in which each lever press (LP) was followed by reward delivery (RD; 100% probability) and, in 50% of trials, a 1-s 5 Hz red laser stimulation (Ls) to optogenetically activate CINs and disrupt the ACh dip–rebound. **B.** Example raster plots of LPs (orange ticks) and magazine entries (MagEnt; blue ticks) in control (no-laser) trials (left) and laser trials (right). **C.** Average distribution of MagEnt relative to LP onset for control (gray) and laser (red) trials (n = 6 mice), showing similar timing of reward collection between conditions. **D.** Heatmaps of z-scored ACh dynamics aligned to RD for control (left) and laser (right) trials. **E.** Trial-averaged ACh dynamics (mean ± SEM) showing that laser stimulation abolished the dip–rebound pattern (n = 9 mice). **F.** Statistics show that the 1-second 5 Hz laser delivery after reward delivery diminished the amplitude of the ACh dip-rebound after reward delivery (Dip: *t*_(8)_ = 3.26, \**p* <0.05, paired t test; Rebound: *t*_(8)_ = 2.77, \**p* <0.05, paired t test).

